# Inhibitors of lysinoalanine crosslinking in the flagella hook as antimicrobials against spirochetes

**DOI:** 10.1101/2024.11.01.621575

**Authors:** Michael J. Lynch, Kurni Kurniyati, Maithili Deshpande, Nyles W. Charon, Chunhao Li, Brian R. Crane

## Abstract

Spirochetes are especially invasive bacteria that are responsible for several human diseases, including Lyme disease, periodontal disease, syphilis and leptospirosis. Spirochetes rely on an unusual form of motility based on periplasmic flagella (PFs) to infect hosts and evade the immune system. The flexible hook of these PFs contains a post-translational modification in the form of a lysinoalanine (Lal) crosslink between adjacent subunits of FlgE, which primarily comprise the hook. Lal crosslinking has since been found in key species across phylum and involves residues that are highly conserved. The requirement of the Lal crosslink for motility of the pathogens *Treponema denticola* (Td) and *Borreliella burgdorferi* (Bb) establish Lal as a potential therapeutic target for the development of anti-microbials. Herein, we present the design, development and application of a NanoLuc-based high-throughput screen that was used to successfully identify two, structurally related Lal crosslink inhibitors (hexachlorophene and triclosan) from a library of clinically approved small molecules. A structure-activity relationship study further expanded the inhibitor set to a third compound (dichlorophene) and each inhibitor was demonstrated to biochemically block autocatalytic crosslinking of FlgE from several pathogenic spirochetes with varied mechanisms and degrees of specificity. The most potent inhibitor, hexachlorophene, alters Lal crosslinking in cultured cells of Td and reduces bacterial motility in swimming plate assays. Overall, these results provide a proof-of-concept for the discovery and development of Lal-crosslink inhibitors to combat spirochete-derived illnesses.

## Introduction

Spirochetes rely on chemotaxis and motility to cause several important human diseases that include Lyme disease (caused by *Borreliella burgdorferi*, Bb), syphilis (*Treponema pallidum* subspecies *pallidum*, Tps), yaws (*Treponema pallidum* subspecies *pertenue*, Tpy), periodontal disease (*Treponema denticola*, Td), and leptospirosis (*Leptospira interrogans*, Li).^1–4^ Lyme disease (LD) is the predominant tick-borne illness in the United States, with the CDC reporting that over 475,000 people were treated for LD in 2022 across an ever-increasing geographical range centered in the Northeastern United States.^5,6^ Spirochetes are highly invasive bacteria, and their unique motility and chemotaxis are essential for virulence.^1,3^ Pathogenic spirochetes generally enter mammalian hosts through either skin or mucous membranes. In contrast to wild-type cells, non-motile and non-chemotactic mutants of Bb fail to disseminate into host tissues on deposition in the skin of mice by either syringe inoculation or tick bite.^7–10^ Spirochete motility uniquely attributes to periplasmic flagella (PFs) that are encased in the periplasmic space (Figure 1A), and thereby shielded from the immune system.^11^ PFs are driven by rotary motors at the base in a manner similar to that of other bacterial flagella; however, in spirochetes the PFs rotate in the periplasm.^1,3^ In species such as Bb, the PFs are also skeletal organelles, e.g. mutants of Bb lacking PFs have markedly altered cell shapes, and as such, are rod-shaped instead of flat-waved.^1,3,12,13^ Rotating PFs exert force on the flexible cell cylinder to generate backward-moving waves that allow the cell to swim in a given direction.^1–3,14,15^ This unique form of motility allows spirochetes to penetrate and transverse in connective tissues such as the collagen matrix of skin that otherwise inhibit the motility of most other bacteria. It also enables these organisms to be highly tissue invasive.^1–3,11,16,17^ Most important, mutants deficient in motility or chemotaxis are markedly less virulent than wild-type (WT) cells.^1,3,8–10,18,19^

**Figure 1:**
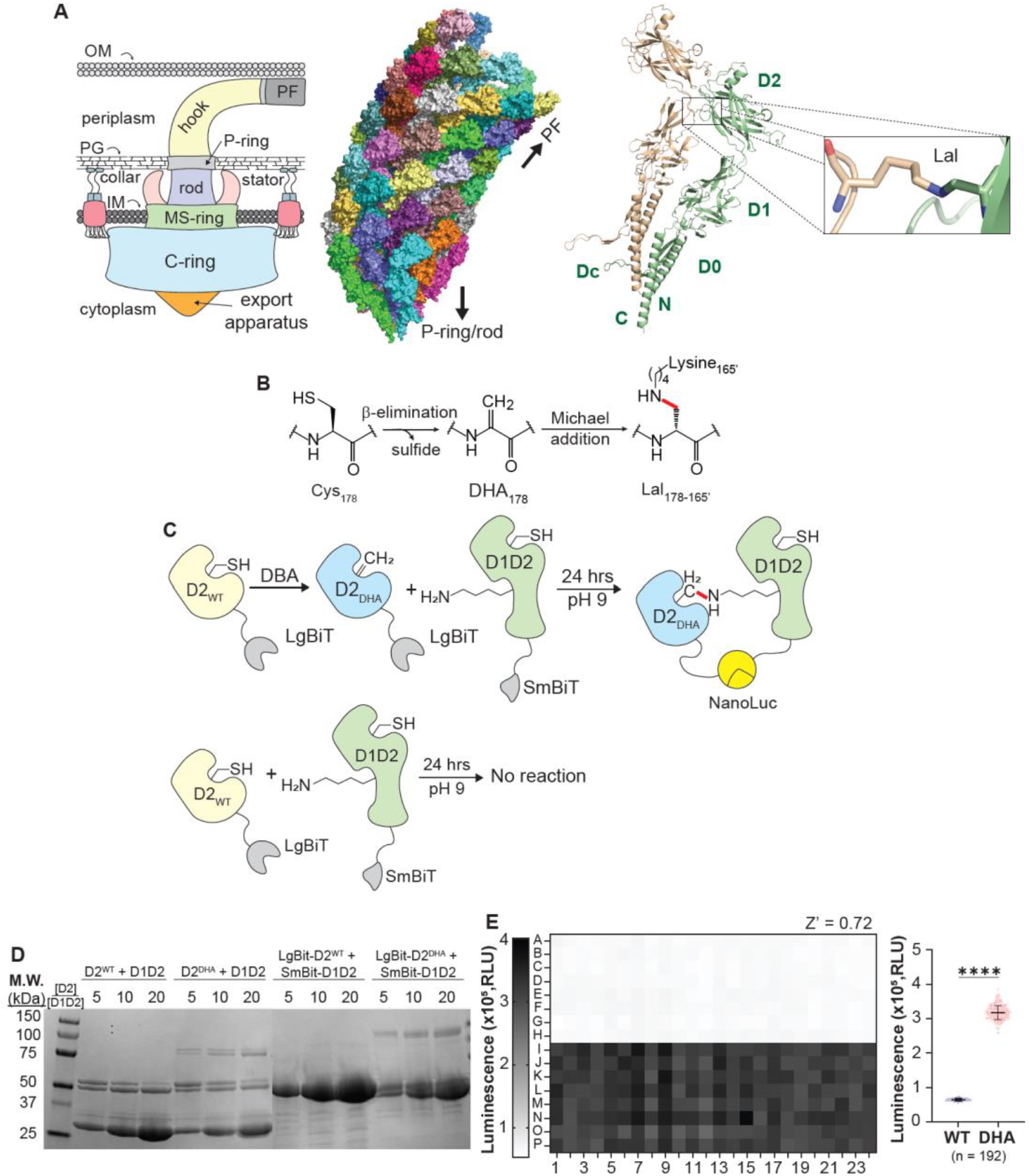
High-throughput screen to identify Lal crosslinking inhibitors. **A)** Schematic of motor components (left), structure of the flagellar hook (PDB 6K3I) ^32^ with individual FlgE subunits colored differently (middle) and location of the Lal crosslink between adjacent FlgE subunits (right). The model of Lal-crosslinked full-length *T. denticola* FlgE (right) was generated from the truncated Lal-crosslinked dimer (PDB 6NDX) ^27^ and AlphaFold-generated FlgE monomer models. **B)** Biochemical mechanism of Lal crosslink formation in *T. denticola* FlgE. **C)** Overview of NanoLuc-based high-throughput screen used to identify FlgE Lal crosslink inhibitors. To create a reactive Td FlgE D2 (Ala168-Thr334) species, the native Cys178 in TdFlgE LgBiT-D2^WT^ was converted to dehydroalanine (DHA, LgBiT-D2^DHA^) via treatment with 2,5-dibromohexadiamide (DBA). LgBiT-D2^DHA^ is then mixed with Td FlgE SmBiT-D1D2 and incubated for 24 hours to afford Lal-crosslinked FlgE heterodimers and a functional NanoLuc enzyme. LgBiT-D2^WT^ serves as a negative control because no Lal crosslinking occurs unless Cys178 is converted to DHA. **D)** SDS-PAGE crosslinking assay of Td FlgE_D1D2_ (D1D2) incubated with increasing concentrations of either wild-type FlgE_D2_ (D2^WT^) or DHA FlgE_D2_ (D2^DHA^). NanoLuc SmBit-D1D2^WT^ and LgBit-D2^WT/DHA^ fusions were also tested for Lal crosslink formation. **E)** Z’ plate of high-throughput screen. (left) Raw chemiluminescence values are represented by a heatmap with negative controls (-Lal) in rows A-H and positive controls (+ Lal) in rows I-P (Z’ = 0.72). (right) Summary of Z’ plate data. The HTS was performed in a 384-well plate with a final volume of 30 μL/well. Statistical significance was calculated using a two-tailed Student’s t-test (p<0.001, ^****^).

At the base of a PF lies a universal joint, known as the hook, that links the motor to the long filament.^2,20,21^ The hook is a well-conserved tubular structure composed of multiple FlgE proteins. FlgE varies in size and sequence among bacteria, and this variation optimizes motility in specific ecological niches.^22–25^ As the hook rotates, its constituent ∼120 FlgE subunits must alter their interactions to compensate for large changes in curvature.^21,26^ The spirochete hook is unique; it must operate in the particularly confined environment of the periplasm, very likely requiring unique stabilization and fortification.^1^ We described a new type of inter-molecular protein crosslink stabilizes the interactions among FlgE subunits of spirochetes in the form of lysinoalanine (Lal), which forms spontaneously between conserved cysteine and lysine residues on adjacent subunits (Figure 1A-B).^27–29^ Natural Lal production in biological systems was previously limited to post-translationally modified antimicrobial peptides, such as the lantibiotics, which requires a suit of enzymes to catalyze the modification.^30,31^ In contrast, we have shown that Lal biosynthesis by spirochete FlgE is autocatalytic and requires no enzymes or cofactors.^27^ In Td, Bb, and Tp, FlgE is composed of four distinct domains: D0, Dc, D1, and D2 (Figure 1A). The Lal cross-link forms between Cys178 of D2 and Lys165 of the D1-D2 linker. In addition to Cys178 and Lys165, Lal crosslinking depends on the invariant residue Asn179 and proceeds through a β-elimination of Cys178 to form sulfide and the intermediate dehydroalanine (DHA, Figure 1B).^27^ Specific hydrogen bonding interactions activate DHA for aza-Michael addition with Lys165.^27^ Our results suggest that all spirochete clades conserve the crosslink and its formation mechanism, although, in Li, the analog of Cys178 is replaced by a Ser residue, which nonetheless is still capable of forming Lal, albeit with less specificity.^28^

Owing to the dependencies of virulence, motility and Lal crosslinking in spirochetes, we sought to develop a high-throughput screen (HTS) for the identification Lal crosslink inhibitors. The optimized HTS is based on the complementation of a split-luciferase enzyme (NanoLuc^33,34^) by the covalent association of two engineered and chemically primed FlgE domains. Upon subjecting the NIH clinical compound library to the HTS two related Lal crosslink inhibitors were identified, with a third arising from a structure-activity relationship (SAR) study. The most potent inhibitor, hexachlorophene, perturbs Lal crosslinking in Td cell culture and commensurately reduces Td motility in swimming plate assays. We herein describe the screening strategy and implementation that provides a path forward for the discovery and development of therapeutic small molecules tailored to the motility characteristics of pathogenetic spirochetes.

## Results and Discussion

### Development of a NanoLuc-based Lal crosslinking HTS

Lal formation requires FlgE subunits to associate with one another via their D0 domains. Removal of these domains results in soluble FlgE fragments that do not interact, thereby halting Lal formation.^27^ We hence created an HTS based on two engineered FlgE truncations: FlgE D2 domain (D2, Ala168-Thr344) and FlgE D1D2 domain (D1D2, Asn91-Ser423) that do not interact with one another and cannot produce a covalent Lal crosslink. D1D2 and D2 were then fused at their C-termini to the NanoLuc split-fusions SmBiT and LgBiT, respectively, to create the SmBiT-D1D2 and LgBiT-D2 fusion proteins, which can be produced in large amounts (Figure 1C). To initiate Lal formation between these proteins, Cys-178 in LgBiT-FlgE_D2_ (LgBiT-D2^WT^) was converted to DHA (LgBiT-D2^DHA^) via pre-treatment with 2,5-dibromohexanediamide (DBA, Figure 1C). Due to the high reactivity of the DHA electrophile^27^, when LgBiT-D2^DHA^ is mixed with SmBiT-D1D2, Lal crosslinking proceeds spontaneously despite the lack of D0 domains on either construct. No Lal crosslinking is observed in a similar reaction where LgBiT-D2^WT^ is used in lieu of LgBiT-D2^DHA^ (Figure 1C and D); an observation we exploited in establishing a negative control for the HTS. To identify Lal inhibitors, NanoLuc enzymatic activity becomes a readout for FlgE protein-protein interactions that depend on the presence of Lal. When no Lal crosslink forms, either because of small-molecule inhibition or when LgBiT-D2^WT^ is used, the two split-fusions do not interact to form active NanoLuc, resulting in low luminescence values upon substrate addition (Figure 1E). Upon Lal formation, SmBiT and LgBiT associate to form a functional NanoLuc that produces stable luminescence for >60 minutes (Figure 1E and Figure S2A).

Practically, the HTS functions in two steps. First, D2^DHA^ and D1D2 are mixed at a 10:1 stoichiometric ratio and incubated for 24 hours to allow for Lal crosslinking. Previous work has shown that increased D2^DHA^:D1D2 ratios drive Lal formation by supplying more of the reactive DHA intermediate. Upon Lal formation, the D1D2:D2 heterodimer positions the SmBiT and LgBiT such that they associate into a functional NanoLuc capable of turning over substrate and producing chemiluminescence. To complete the assay, equal parts of a 1:1000x dilution of Nanluc substrate (Fz) are added to each well and the luminescence measured. The positive control for this assay was a 10:1 mixture of D2^DHA^ and D1D2, whereas the negative control was D2^WT^ and D1D2. The vehicle DMSO also served as a carrier control. The assays functioned well on a 384-well scale, producing a Z’ of 0.72 during initial HTS testing (Figure 1E) and an average Z’ of 0.76 ± 0.06 for all compound plates screened.

### Identification of compounds that alter Lal crosslinking

The overall screening platform had four distinct phases (Figure 2A): phase 1 - NanoLuc HTS, phase 2 - NanoLuc control test, phase 3 – secondary crosslinking assays by SDS-PAGE and phase 4 - *T*.*denticola* cell culture studies (Figure 2A). In phase one, we tested compounds from the NIH clinical collection (NIHCC) in duplicate and each compound produced results that segregated into one of three categories: compounds with chemiluminescent values greater than the average values of the positive controls plus five times their error (+5σ_p_), (2) compounds with less than the values of the positive controls minus three times their error (-3σ_p_), and (3) compounds with values between +5σ_p_ and -3σ_p_ (Figure 2B). Compounds in the first two categories that produced consistent replicates progressed to the next phase of the workflow.

**Figure 2:**
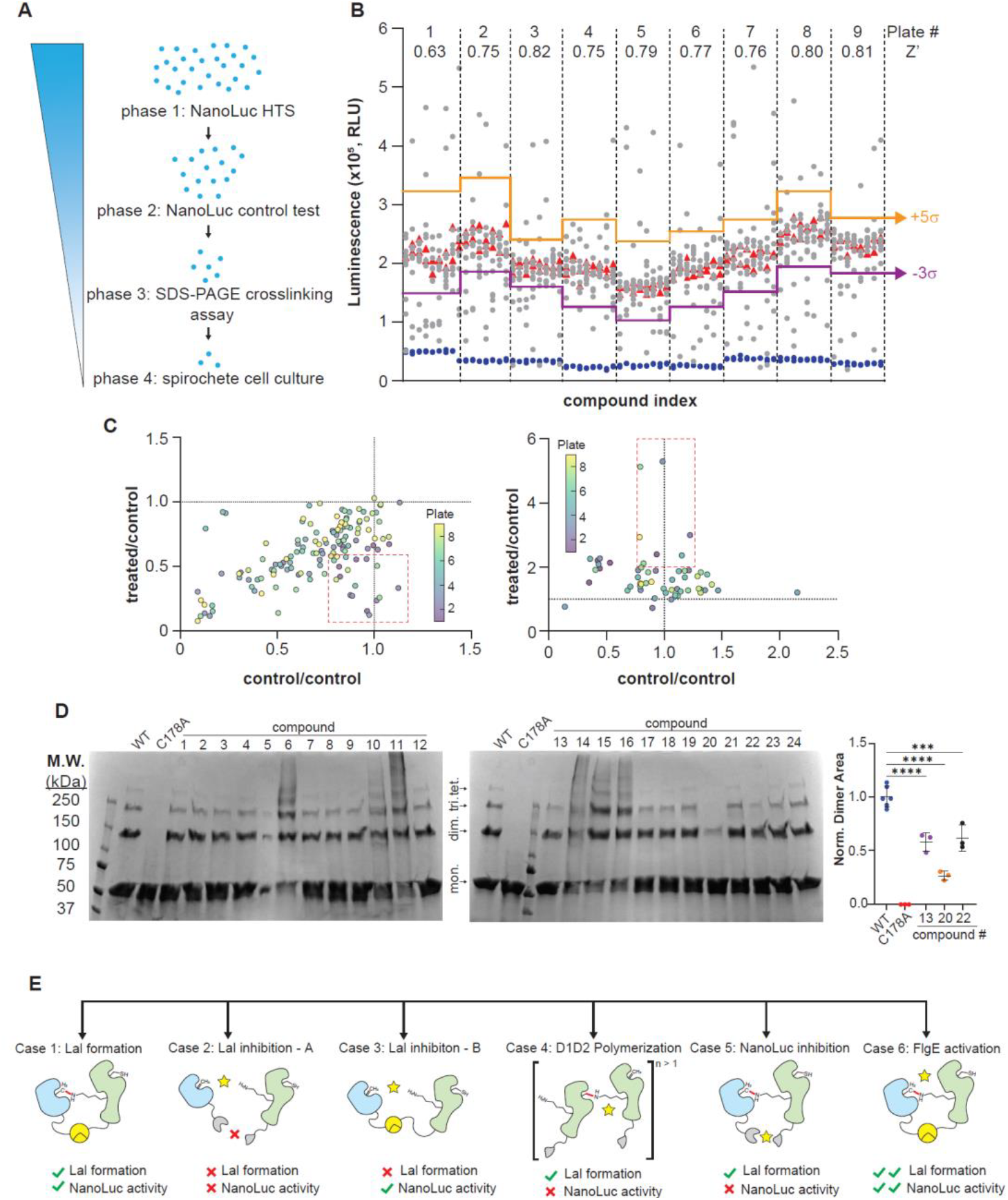
Identification of Lal-crosslink modulating compounds. **A)** Overview of inhibitor discovery. Compounds were screened first using the NanoLuc HTS *(phase 1)* to find hits that are then re-screened to control for NanoLuc interference *(phase 2)*. Candidate compounds are then tested in SDS-PAGE crosslinking assays with unmodified WT FlgE *(phase 3)*. Top ranked compounds were then tested on *T. denticola* cell cultures for Lal crosslinked FlgE subunits and cell motility *(phase 4)*. **B)** Screening results from the NIHCC. For each plate, the Z’ value, positive control (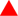), negative control (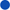), +5σ (upper orange line) and -3σ (lower purple line) values are included. Compounds with chemiluminescence values ≤ 3σ and ≥ 5σ were tested in step 2. **C)** NanoLuc control assay from compounds with chemiluminescence values ≤ 3σ (top) and ≥ 5σ (bottom) identified in step 1 (x-axis: RLU^0hr^/RLU^DMSO^ ratio, y-axis: RLU^24hr^/RLU^DMSO^; RLU, relative luminescence units). Compounds within the red dotted boxes (n=24) have RLU^0hr^/RLU^DMSO^ values = 1 ± 20%, with RLU^24hr^/RLU^DMSO^ ≤ 0.5 (left) or ≥ 2 (right) and were selected for step 3. **D)** SDS-PAGE crosslinking assay of Td FlgE_FL_ (20 μM) incubated with compound (333 μM). Lal-crosslinked proteins band intensities were compared to WT Td FlgE_FL_ (positive control) and C178A Td FlgE_FL_ (negative control). Top candidate inhibitors were measured in triplicate and the dimer band intensity plotted. See Figure S5A for SDS-PAGE gels. Statistical significance was calculated using a one-way ANOVA (p<0.05^*^, 0.01^**^ and 0.001^****^). **E)** Different outcomes observed in our NanoLuc HTS after SDS-PAGE assay (see text).

In phase 2, the phase 1 compounds were evaluated for direct effects on NanoLuc activity by taking advantage of the relatively slow rate of Lal crosslinking. Compounds were added to wells containing D2^DHA^ and D1D2 at 0-hour and 24-hour timepoints and the luminescence measured and compared. Control wells (DMSO only) were also included for comparison. If a compound inhibited NanoLuc activity directly, luminescence was decreased equally at 0-hours or after 24-hours. In contrast, Lal inhibitors required the 24-hour incubation to show a decrease in luminescence. To quantitatively identify hit compounds, scatter plots were generated where the 24-hour/DMSO-only luminescence ratio (treated/control, y-axis: RLU^24hr^/RLU^DMSO^) were plotted against the 24-hour/0-hour time point luminescence ratio (control/control, x-axis: RLU^0hr^/RLU^DMSO^) for each compound (Figure 2C). DMSO-only controls yielded an average standard deviation of approximately 7%, thus a 20% (∼3σ) control/control window was used to filter compounds in the x-axis. Compounds in the y-axis (treated/control) were filtered differently depending on if they decreased (inhibitors) or increased (activators) luminescence compared to the DMSO-only controls. Compounds were considered inhibitors only if they had luminescence values equal to or less than 50% compared to the DMSO-only control. Compounds were considered activators if they produced luminescence values greater than 2-fold higher than DMSO-only controls. Compounds within these values were tested further in step three of our screen and the remaining compounds were excluded.

Previously, we developed an SDS-PAGE Lal crosslinking assay that was used in mutagenesis experiments to identify residues of FlgE required for Lal formation.^27^ This assay employs SDS-PAGE to monitor the formation of Lal-crosslinked high-molecular weight complexes (HMWCs) comprised of unmodified, recombinantly produced, full-length Td FlgE (FlgE_FL_). In phase three, we tested compounds by SDS-PAGE to rule out the possibility of compounds only inhibiting Lal crosslink formation in the modified forms of FlgE used for the HTS (DHA or truncated) but not in the wild-type (WT) FlgE_FL_. Phase two identified 24 compounds that either increased or decreased the luminescence signals compared to the controls. At this time, we hypothesized that increased signals may represent compounds that activate the crosslinking chemistry. For example, the cysteine-reactive compounds N-ethylmaleimide (NEM) and Ellman’s reagent (DTNB) activate Lal formation in FlgE_FL_ by favoring elimination of sulfide from a Cys178 adduct.^27^ The NanoLuc HTS utilizes the DHA form of the D2 domain, thus any compound that activates Lal formation in the HTS assay must be doing so via a different mechanism. Regardless, to test the 24 compounds, we incubated each compound with WT Td FlgE_FL_ in crosslinking buffer for 48 hours. During this time, FlgE_FL_ will self-assemble, and Lal crosslink formation will follow accordingly. The protein composition of each sample was assessed via SDS-PAGE and compared to WT (DMSO only) and Cys178Ala (C178A) Td FlgE_FL_ as positive and negative controls, respectively (Figure 2D). From the gel assay, compounds 6, 10, 11, 14, 15, 16 and 31 (Figure 2D and Figure S4) appeared to increase the amount of Lal-crosslinked HMWCs, while compounds 13, 20, and 22 reduced Lal crosslinking. Compound 5 also initially appeared to reduce Lal crosslinking; however, the effect was not reproducible. Compounds that increased Lal crosslinking (i.e. activators) were investigated in additional experiments (see below). Compounds that decreased Lal crosslinking (i.e. inhibitors) were tested in triplicate and the Lal crosslinking quantified and significance measured (Figure 2D and Figure S5A). Hit compounds were also evaluated in SDS-PAGE Lal crosslinking using the D2^DHA^ and D1D2 NanoLuc split-fusions (Figure S4A). These assays support the findings presented in Figure 2D,thereby validating the ability of the compounds to inhibit Lal crosslinking in WT FlgE and the screened truncations.

Together, the NanoLuc HTS and SDS-PAGE assays segregate hit compounds into one of two categories: compounds that (1) increase or (2) decrease Lal crosslink formation; however, when considering the potential outcomes of our NanoLuc HTS, there are additional scenarios that may produce perturbed luminescence values (Figure 2E). For example, a compound may produce at least six potentials outcomes: case 1 – compound has no effect on Lal crosslink formation and Nanoluc activity is normal (the majority case represented by a non-interacting compound), case 2 – compound inhibits Lal formation and no Nanoluc activity is observed (a bona fide inhibitor), case 3 – Lal crosslinking is inhibited but Nanoluc activity is observed nonetheless (such inhibitors will be missed by the assay), case 4 – Lal crosslinking occurs between the SmBiT-FlgE_D1D2_ fusions due to activation of the Cys178, but no NanoLuc activity is observed (such activators will be missed by the assay), case 5 – compound inhibits NanoLuc but not Lal crosslinking (controlled for by differentiating 0 hr vs 24 hr time points) and case 6 – compound drives Lal crosslink formation at levels higher than the positive controls, leading to increased NanoLuc activity (a bona fide activator).

### Top ranked compounds inhibit Lal crosslinking in FlgE orthologs from diverse spirochete species

The FlgE inhibitor screening workflow successfully identified two compounds (20 and 22) from a library of 727 compounds. These compounds, known as hexachlorophene (20, HC) and triclosan (22, TC) are used as antiseptics and applied topically.^35,36^ Overall, the structures of HC and TC are similar; consisting of two polychlorinated phenol rings fused via a bridging methyl- or oxo-group, respectively (Figure 3A). Lal crosslinking is a conserved post-translational modification catalyzed by FlgE orthologs across the spirochete phylum. We thus tested the inhibitory properties of HC and TC on other, more distantly related orthologs of *T. denticola* FlgE_FL_ with the SDS-PAGE crosslinking assay. Both HC and TC inhibit Lal crosslinking by FlgE from *B. burgdorferi, T. pallidum* and *L. interrogans*, albeit with varying effectiveness (Figure 3A).

**Figure 3:**
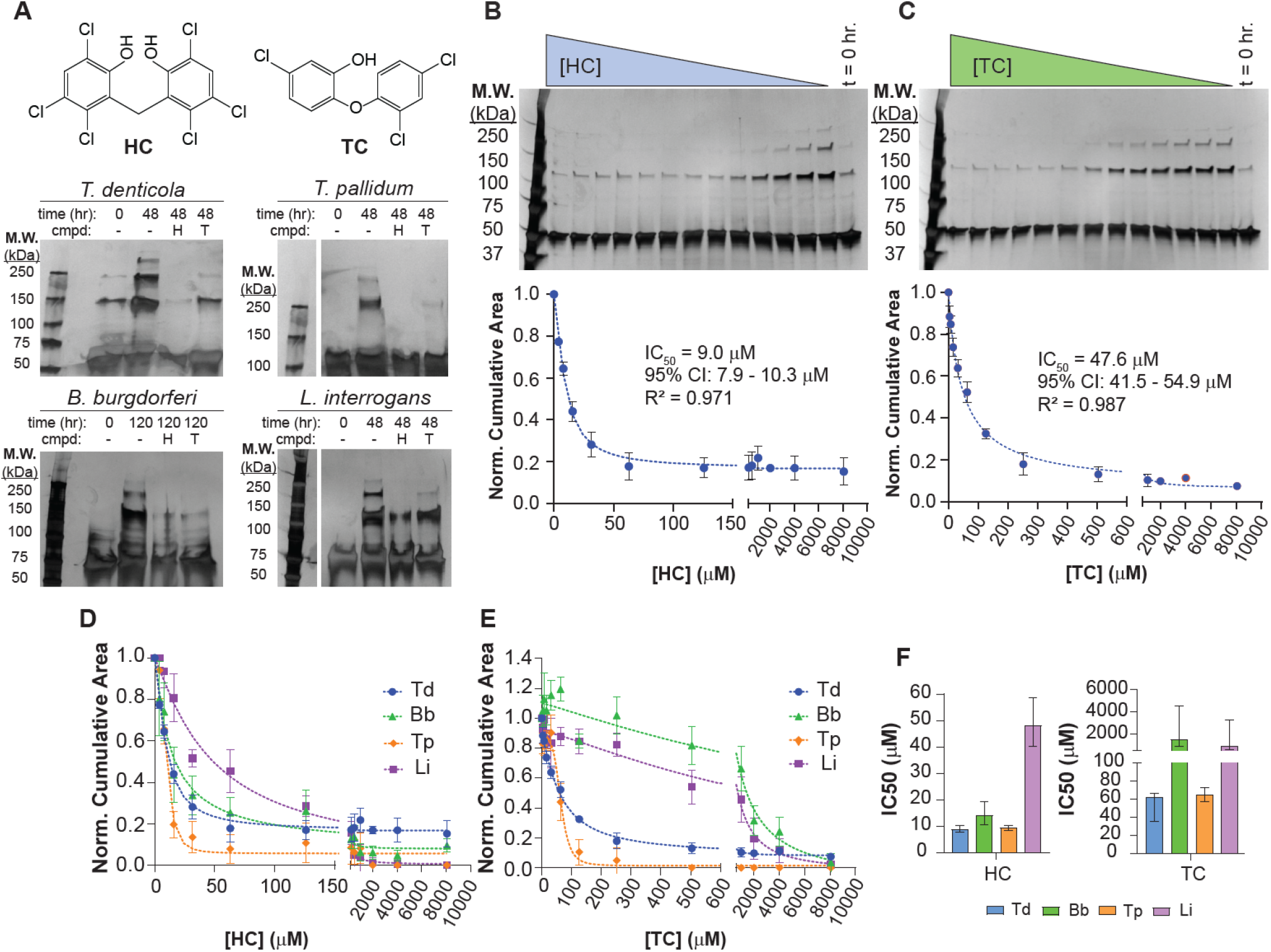
Characterization of candidate Lal crosslink inhibitors. **A)** SDS-PAGE gels of crosslinking assays of 20 μM FlgE orthologs from *T. denticola* (Td), *T. pallidum* (Tp), *B. burgdorferi* (Bb) and *L. interrogans* (Li) with 1 mM HC or TC. SDS-PAGE IC_50_ determination of **B)** HC and **C)** TC for Td FlgE Lal crosslinking. Replicate IC_50_ curves for **D)** HC and **E)** TC for FlgE orthologs. Normalized cumulative band areas are presented as the average area ± the standard deviation of three replicates. **F)** IC_50_ values for (left) HC and (right) for Td, Tp, Bb and Li FlgE determined from the data shown in (D) and (E). IC_50_ errors reported as the error from the fit (Figure S6).

### IC50 determination of HC and TC

We next sought to determine the IC_50_ for both HC and TC against Td FlgE_FL_ with the *in vitro* SDS-PAGE crosslinking assay. HC and TC were titrated from concentrations ranging from 8 μM to 8 mM and the cumulative Lal-crosslinked band area measured on silver-stained SDS-PAGE gels (Figures 3B and C). Band area densities were measured in triplicate (Figure S5B) and summed for all observable oligomer bands, normalized against DMSO only levels and the data fit to a four-parameter [inhibitor] vs. response equation (Figure S6) to determine the IC_50_ values for each compound. Overall, HC is a more potent inhibitor with an IC_50_ of 9 μM (95% C.I. = 7.9 μM – 10.3 μM) compared to TC with an IC_50_ of 47.6 μM (95% C.I. = 41.5 μM – 54.9 μM). IC_50_ curves for HC (Figure 3D) and TC (Figure 3E) show similar trends with Bb, Tp and Li FlgE that were observed for Td FlgE (Figure 3B-C) with some minor differences. HC exhibits similar potencies against Bb and Tp FlgE compared to Td FlgE, with IC_50_’s of 14.2 μM (95% C.I. = 10.7 μM – 19.3 μM) and 9.37 μM (95% C.I. = 8.44 μM – 10.5 μM), respectively (Figure 3F and Figure S6). In contrast, HC is a less potent inhibitor of Lal crosslinking for Li FlgE, with an IC_50_ of 48.3 μM (95% C.I. = 40.2 μM – 58.6 μM). TC inhibits Lal crosslinking in Tp FlgE at levels comparable to Td FlgE, having an IC_50_ of 64.7 μM (95% C.I. = 56.9 μM – 72.4 μM), whereas TC weakly inhibits Bb and Li FlgE, with IC_50_’s of 1.101 mM (95% C.I. = 1.018 mM –1.154 mM) and 909 μM (95% C.I. = 879 μM – 1.013), respectively (Figure 3F and Figure S6).

### HC and TC inhibit Lal crosslinking via different mechanisms

Given that the HTS targeted the reactive DHA intermediate in Lal formation, HC and TC could potentially form adducts with these species and act as covalent inhibitors. Thus, we investigated the reversibility of inhibition by these compounds using SDS-PAGE Lal crosslinking assays. D2^DHA^ was treated with DMSO, HC, TC, or β-mercaptoethanol (βME) and then extensively dialyzed to remove any non-covalently bound compounds. βME served as a covalent inhibitor control because it is known to react with DHA via a thia-Michael reaction to form a βME-adduct.^27^ Treated and dialyzed samples were then incubated with D1D2 and the formation of D2:D1D2 heterodimers was monitored via SDS-PAGE. DMSO- and TC-treated D2^DHA^ samples were still able to form Lal-crosslinked heterodimers at levels comparable to the untreated control (Figure 4A). HC- and βME-treated samples do not form Lal crosslinked heterodimers after dialysis, thereby indicating that HC may be a covalent inhibitor, whereas TC is a non-covalent inhibitor. These mechanistic differences hold for WT FlgE_FL_, (Figure S7A). For reasons unknown, βME appears to have both covalent and non-covalent inhibition properties, depending on whether truncated or non-truncated FlgE proteins are used. The rate of Lal inhibition by HC was evaluated by performing FlgE crosslinking after dialysis following different times of HC treatment. By this metric, HC inhibition is quite rapid, with nearly 100% of the inhibition produced after only a few minutes of HC incubation (Figure S7B).

**Figure 4:**
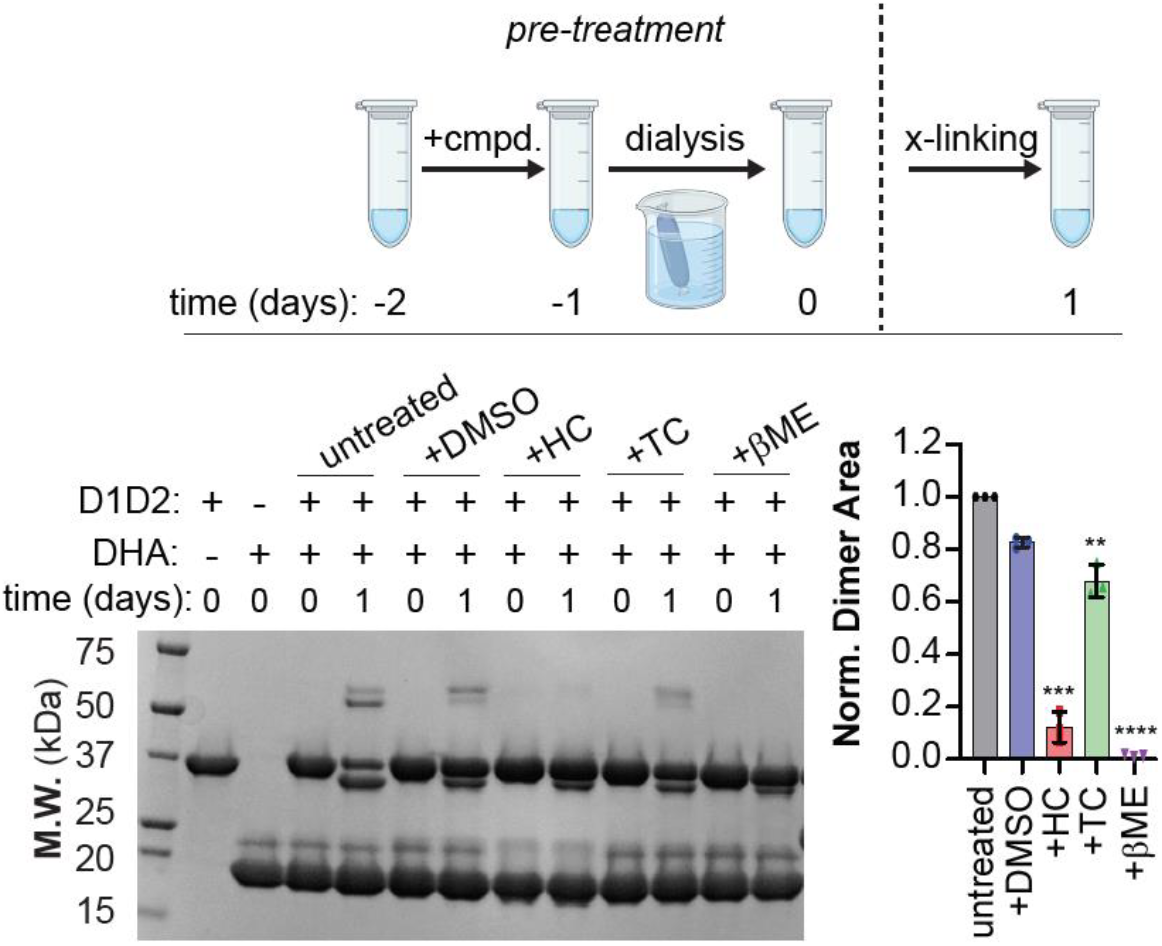
Differential inhibition by HC and TC. **A)** SDS-PAGE Lal crosslinking assay with 20 μM D2^DHA^ incubated with 1 mM HC, TC or 100 mM βME. (top) Overview of experimental design of assay. Each sample was treated with DMSO or compound for 24 hours at 4°C. Samples were then dialyzed to remove any non-covalently attached compound and mixed with D1D2 in a 5:1 D2:D1D2 ratio. Samples were incubated for 24 hours and crosslinking was monitored via SDS-PAGE. (bottom - left) SDS-PAGE gel of compound-treated samples post-dialysis. For this assay, the untreated (no compound, DMSO or dialysis) and DMSO-treated samples serve as positive controls. βME reacts with DHA via a thia-Michael addition and is covalent inhibitor. (bottom - right) Quantification of Lal-crosslinked FlgE protein oligomer bands. Total band areas were normalized to untreated sample levels and reported as the average ± the standard deviation of three replicates. Statistical significance was calculated using a one-way ANOVA comparing DMSO with HC, TC and βME-treated samples (p<0.05^*^, 0.01^**^ and 0.001^****^).

**Scheme 1:**
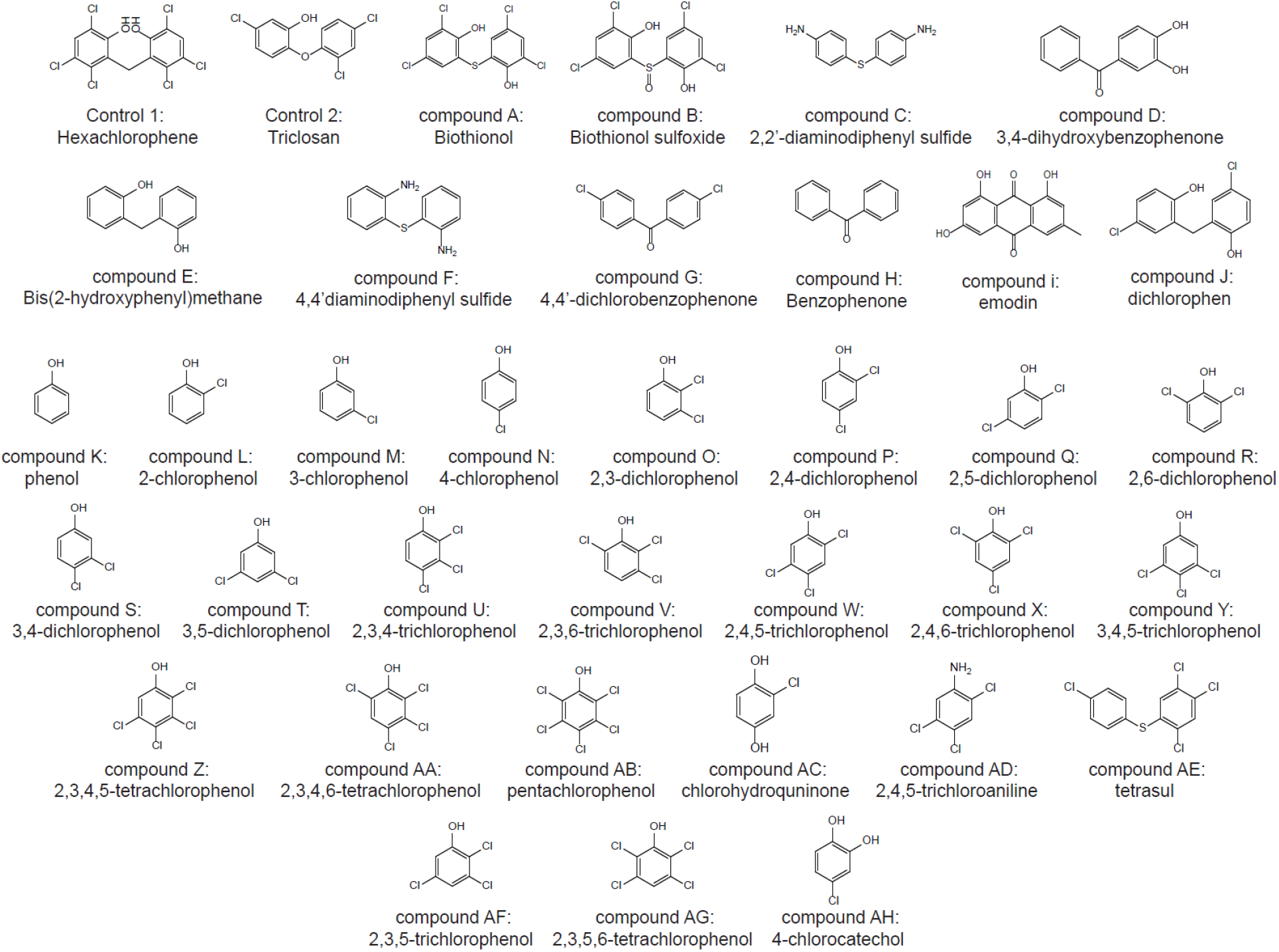
Compounds tested for structure-activity relationship studies

### Structure-activity relationship (SAR) studies of HC and TC

Thirty-four compounds with structures that share common features with one or both of HC and TC were tested as Lal inhibitors (Scheme 1). The compounds tested consisted of several commercially available polychlorinated bis-phenols (bithionol, biothionol sulfoxide and dichlorophen). These compounds vary in the overall relative location and position of the chloro-groups on the rings, as well as bridging group identity and hybridization. We also explored fused-ring systems without hydroxyl groups (2,2-diaminodiphenyl sulfide, 4,4’diaminodiphenyl sulfide, 4,4’-dichlorobenzophenone, benzophenone and tetrasul), phenolic-derivatives lacking chloro-substituents (3,4-dihydroxybenzophenone and Bis-(2-hydroxyphenl)-methane), as well as a more distantly related planar, three-ring phenolic-derivative (emodin). We also explored single-ring polychlorinated phenols to determine if both rings were required and if one of the rings in HC or TC was more important than the other for inhibition of Lal crosslinking. These compounds consisted of all mono-, di-, tri-, tetra- and penta-substituted chlorinated phenols commercially available, as well as other, more distantly related compounds (chlorohydroquinone, 2,4,5-trichloroaniline and 4-chlorocatechol). We tested each compound in the SDS-PAGE crosslinking assays compared to DMSO-only positive controls (Figure S8). If compounds appeared to inhibit Lal formation, we then performed the assay again but varied the concentration of compound to establish a dose-response effect. Upon observing a dose response, titrations were then performed and the relative IC_50_ value determined from data fitting. Out of the 34 compounds tested, only emodin, bithionol sulfoxide and dichlorophen (DC) had a dose-response effect on Lal crosslinking. Of these three compounds, only dichlorophen altered Lal crosslinking similar to HC and TC, whereas inhibition by bithionol sulfoxide and emodin was very modest. The IC_50_ value for DC was measured to be 308 μM (95% C.I. = 211 μM – 527 μM), making it the weakest of the three inhibitors identified in our screen (Figure S9).

In addition to inhibitors, we also identified several compounds that appeared to activate FlgE and increase Lal crosslinking relative to the positive controls (Figure 2D and Figure S4). These compounds included dopamine (Dopa) and several dopamine-related catecholamines, as well as the cephalosporin-related antibiotic cetirizine (Cef). Interestingly, the catecholamines identified in our assay appeared to cause non-specific crosslinking via either case 4-related or unknown mechanisms (Figure 2E and Figure S4). Of these compounds, Dopa appeared to be the most potent catecholamine tested, forming high-molecular weight complexes (HMWCs) on SDS-PAGE gels even when incubated with protein mixtures that should only be able to form dimers (i.e. D2^DHA^ and D1D2, Figure S4A) and producing the most Lal-crosslinked HMWCs with Td FlgE_FL_ (Figure S4B). These aggregation effects may result from non-specific reactivity between FlgE and catecholamines because this class of compounds are among a major class of Pan-Assay Interference Compounds (PAINS).^37^ This hypothesis was supported by apparent activation by chlorohydroquinone (CHQ) and 4-chlorocatechol (4CC) observed in SAR studies (Figure S8). We performed several SDS-PAGE assays to elucidate if non-specific aggregation was masking activation of Lal crosslinking by these compounds. First, we tested Dopa, CHQ, 4CC and Cef on their ability to affect crosslinking of one-component samples of SmBiT-FlgE_D1D2_ (contains both Cys178 and Lys165), LgBiT-D2^WT^ (contains Cys178, but no Lys165) or LgBiT-D2^DHA^ (contains DHA178, but no Lys165). If these compounds activate FlgE crosslinking, they should have no effect on the D2 proteins alone but may drive Lal crosslinking in D1D2 alone. Comparisons between the DHA or Cys178 forms of LgBiT-D2 evaluate if the activators selectively react with one of these FlgE D2 forms over the other. Two compounds known to facilitate elimination of Cys178 and accelerate crosslinking, N-ethylmaleimide (NEM) and Ellman’s reagent (DTNB), were evaluated as controls. Overall, this experiment revealed that Dopa does produce some non-specific protein crosslinking in all samples tested. Other compounds are unremarkable in their HMWC formation, except for weak dimer band formation in DTNB-treated SmBiT-D1D2 and some weak oligomer bands in all samples treated with CHQ and 4CC (Figure S10A). To further explore the Dopa effect, we performed a titration using the NanoLuc split-fusions (Figure S10B) and Td FlgE_FL_ (Figure S10C). Overall, Dopa gives a dose-dependent effect across all samples tested, which is consistent with both non-specific protein aggregation and Lal activation. The question of aggregation was further addressed by performing the same assay using crosslinking deficient FlgE_FL_ variants Cys178Ala (C178A) and Lys165Ala (K165A), at two different FlgE concentrations and activator ratios: 1) 2 μM FlgE_FL_ and 1 mM activator (activator:FlgE ratio of 500, Figure S10D), and 2) 20 μM FlgE and 333 μM activator (activator:FlgE ratio of ∼17, Figure S10E). Comparing the protein band pattern between both experiments reveals several key findings. First, at high activator:FlgE ratios, Dopa, CHQ and 4CC react indiscriminately with WT, C178A and K165A Td FlgE_FL_; however, the banding pattern for Dopa-treated WT FlgE suggests that only Dopa enhances Lal crosslinking. However, at low activator:FlgE ratios, all compounds produce a crosslinking ladder with WT FlgE_FL_ but not with the variants, except for CHQ, which still shows primarily aggregation. Lastly, compared to NEM- and DTNB-treated Td FlgE_FL_, Cef appears to be a true Lal crosslink activator, because HMSC formation is only observed in WT FlgE_FL_ not in the C178A and K165A variants, with little aggregation at either FlgE concentration (Figure S10D-E).

### HC affects FlgE oligomers in cells and disrupts cell motility

HC and TC have pronounced effects on *T. denticola* cell culture, with HC altering motility and FlgE crosslinking patterns. Previously, we showed that the presence of the Lal crosslink between adjacent FlgE subunits, although not required for proper hook assembly, is required for motility of both *T. denticola* and *B. burgdorferi*.^27–29^ At low concentrations *T. denticola* can tolerate both HC and TC (Figure S11A). HC and TC have antimicrobial properties; thus, the concentrations of both compounds were titrated such that cell growth was minimally affected. At culture concentrations of 83 μM and lower, cell division is only slightly impaired for each compound (Figure S11A). To test if HC or TC influence FlgE oligomers in cells, *T. denticola* were co-cultured with 83.3 μM or 8.3 μM HC (Figure S11B) or TC (Figure S11C) and samples taken at mid-log and stationary phase. For each sample type, the total protein content of the cell lysate was analyzed via stain-free SDS-PAGE and FlgE visualized via αFlgE western-blot. Interestingly, a ∼70 kDa αFlgE-reactive band appeared in stationary-phase samples that were co-cultured with HC. This band, although not at the expected 50 kDa molecular weight of monomeric FlgE, was dose-dependent with HC and was not present in the DMSO-only or ΔFlgE knockout control samples (Figure S11B). This band is absent from TC co-cultured samples, regardless of the growth phase or concentration (Figure S11C), possibly owing to the lower potency of TC compared to HC as reflected by higher IC_50_ measurements. To assess the motility of HC-treated *T. denticola* cells, *T. denticola* cells were passaged for four generations with 83.3 μM HC and ∼150 cells plated on 0.75% (v/v) TYGVS plates supplemented with 83.3 μM HC. Colonies were then picked, grown to stationary phase in the presence of HC and analyzed via SDS-PAGE and western blot. The motility of the bacteria from two colonies were assessed via swimming plate assays and compared to DMSO-only and a non-motile ΔTap1 strains. A ∼70kDa αFlgE-reactive band following HC treatment was replicated in all five of the *T. denticola* colonies tested (Figure 5A). The apparent molecular weight of the new FlgE reactive band that is intermediate to that of a monomer or a dimer may reflect the fact that spirochete flagella proteins are known to undergo additional post-translational modifications when incorporated into PFs.^38^ Moreover, another slightly higher-molecular weight band at ∼85 kDa was also observed in HC-treated samples but not in DMSO-treated controls. The additional band may reflect the additional passage time and growth format of these cultures compared to those shown in Figure S11. The motility of HC-treated cells was also greatly reduced compared to the DMSO-only treated strain (Figure 5B-C). DMSO-treated *T. denticola* cells had a swimming ring diameter of 25 mm, while the two HC-treated colonies had diameters of 8-10 mm, comparable to the non-motile ΔTap1 strain with a diameter of 6 mm. Note that each FlgE subunit crosslinks to two other subunits in the hook and thus a substantial decrease in crosslinking may not produce many monomeric subunits, while nevertheless still compromising the mechanical properties of the hook needed for optimal motility. Overall, these results suggest that HC treatment reduces the size of FlgE oligomers in a time and concentration dependent manner and importantly impairs cell motility.

**Figure 5:**
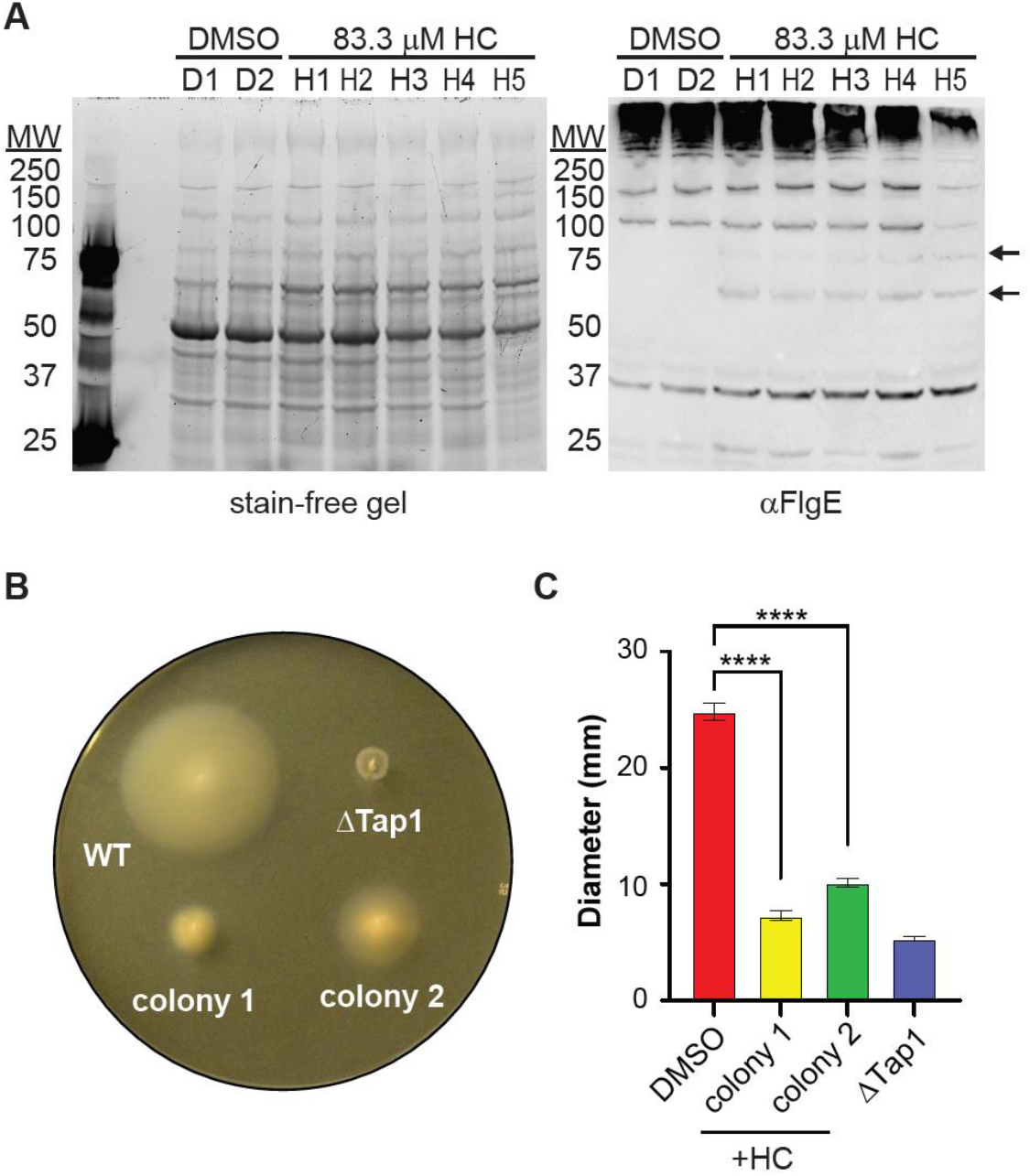
HC alters FlgE in cells and disrupts cell motility. **A)** Stain-free SDS-PAGE gel (left) and αFlgE western blot (right) of different *T. denticola* colonies treated with DMSO (D1-2) or HC (83.3 μM, H1-H5). Black arrows highlight the appearance of a ∼80 kDa and ∼70 kDa band present in HC-treated samples but are lacking in DMSO-treated samples. **B)**Swimming plate assays of *T. denticola* cells under varying treatment (WT = DMSO treatment, ΔTap1 (ΔFliK) = DMSO treatment, non-motile and unflagellated control, colony 1 and colony 2 = 83.3 μM HC treatment). **C**) Swimming plate assays of WT, ΔTap1 and HC-treated *T. denticola* cells. Swimming plate assays were performed in triplicate and the diameter of each strain reported as the average diameter ± the standard deviation of three biological replicates. Statistical significance was calculated using a two-tailed Student’s t-test (p<0.05^*^, 0.01^**^ and 0.001^****^).

### Specific interactions that govern HC inhibition of Lal

We applied computational docking studies to gain a preliminary understanding of how HC interacts with FlgE. Two docking models were generated based on our current hypothesis of HC inhibition. HC inhibition of FlgE crosslinking is either covalent or associated with a very slow off rate. In the former case, HC may modify FlgE via a mechanism not dissimilar to the Lal crosslinking itself. Instead of a 1,4 aza-Michael addition of Lys165 to DHA to yield Lal, HC may react with DHA via a 1,4 oxa-Michael addition. Deprotonation of a phenol group (pKa ∼5) in HC yields a phenolate that could act as a nucleophile and compete with Lys165 for DHA (Figure 6A).^39^ Although oxa-Michael additions have not been heavily utilized in organic synthesis compared to other Michael additions (i.e. aza, thia) due to their notable reversibility under basic conditions at higher temperatures and lower yields, phenol-based alcohols are often more reactive than their aliphatic counterparts.^40^ HC likely intercepts the DHA form of FlgE because the HTS was designed to target DHA and inhibition is only observed under conditions that promote the conversion of Cys178 to DHA (Figure S12). Thus, for the *in silico* docking studies we prepared both a mixed Cys/DHA FlgE_FL_ dimer model (Figure 6B) and a FlgE D2 DHA monomer model based on crystal structures of the various components and intermediates (Figure S13). HC in the phenolate form was docked to each model within a 12×12×12 Å^3^ box centered on the DHA residue. For the FlgE dimer model, top ranked poses place HC within the DHA binding pocket at the interface between two FlgE subunits. Within the cationic DHA pocket, a 3.0 Å hydrogen bond forms between the ε-nitrogen Lys336 and the phenolate oxygen of HC (Figure 6A right panel, docking score = -4.617) and several interactions are made with Ile-162, Asn324, Ala-177, Thr334 (Figure 6B). The phenolate group of HC orients within 3.9 Å of the DHA Cβ, lending support to the oxa-Michael addition mechanism. Docking HC to the D2^DHA^ monomer model produces similar results as the dimer model, with the HC phenolate oxygen again forming a 3.0 Å hydrogen bond between to ε-nitrogen Lys-336 (Figure S13).

**Figure 6:**
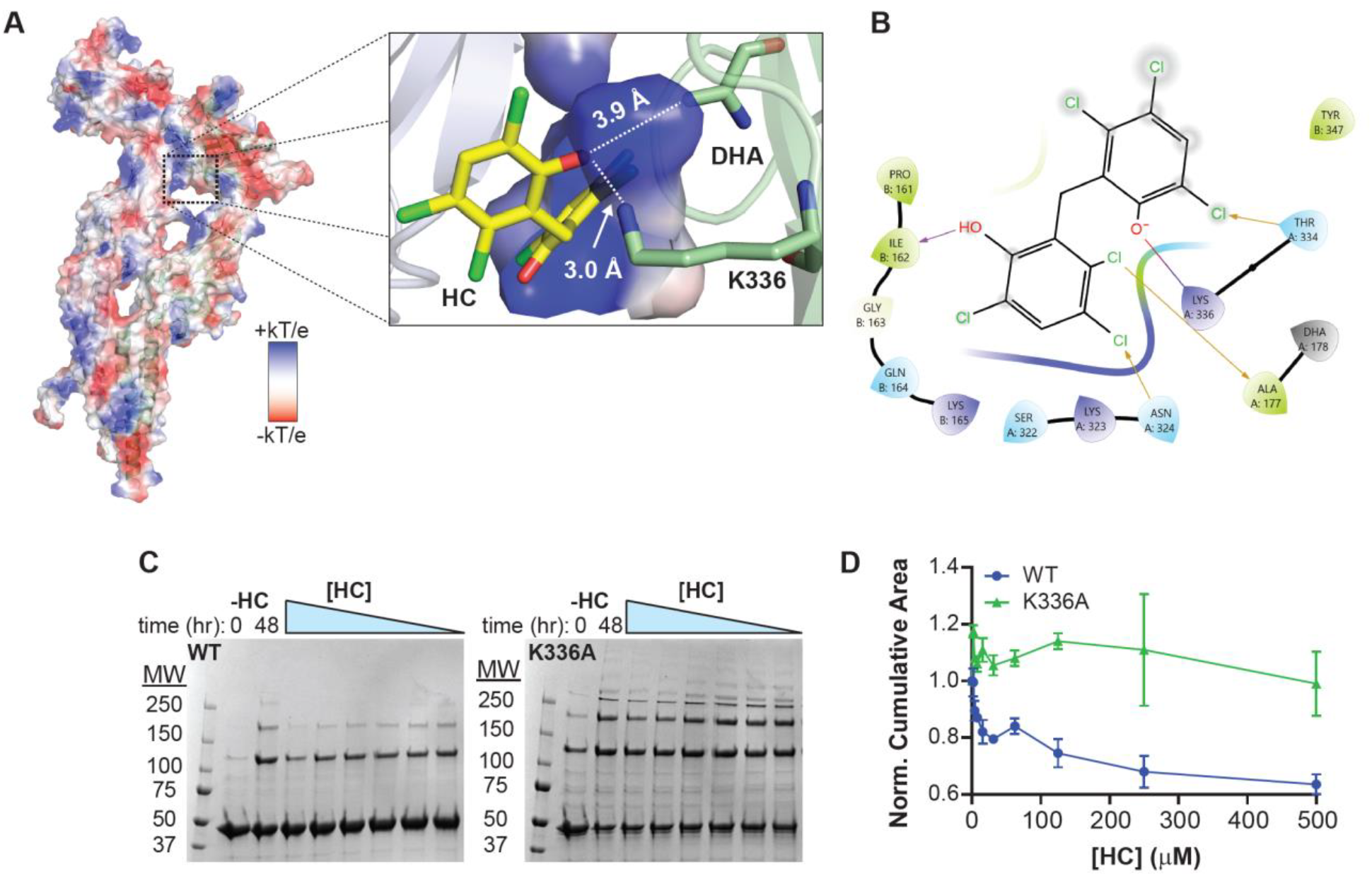
Model of Lal crosslink inhibition by HC. **A)** Top docking pose of HC-bound FlgE (docking score = -4.617). HC was docked to the DHA pocket of a mixed Cys/DHA FlgE dimer model (left, denoted by black dotted box). HC is accommodated well within the DHA binding pocket with a 3.0 Å hydrogen bond to Lys336 and the phenolate within 3.9 Å of the DHA Cβ. **B)** Ligand-interaction diagram between the top docking pose of HC and the FlgE_FL_ dimer model. **C)** SDS-PAGE crosslinking assays with WT (left) and K336A (right) Td FlgE_FL_. For each sample, 20 μM FlgE was incubated with increasing concentrations of HC with Lal crosslinking monitored via SDS-PAGE and stained with Coomassie blue. Each experiment was repeated in duplicate. **D)** Normalized cumulative Lal-crosslinked band areas of WT (blue spheres) and K336A (green triangles) FlgE_FL_ versus [HC] titrations shown in (C). Unlike WT Td FlgE_FL_, whose Lal crosslinked HMWC formation is reduced by ∼40% in the presence of 500 μM HC, Td FlgE_FL_ K336A mutant does not show any reduced Lal crosslink formation, even at HC concentrations exceeding 1 mM.

To test the involvement of the Lys336 side chain in HC-mediated Lal crosslink inhibition, we measured concentration dependent HC inhibition of Lal crosslinking in WT and a Lys336Ala (K336A) variants of Td FlgE_FL_. Unlike WT Td FlgE_FL_, whose Lal crosslinked HMWC formation is reduced substantially by HC, the Td FlgE_FL_ K336A variant does not show any reduced Lal crosslink formation, even at HC concentrations exceeding 1 mM. These results lend support to the docking results that indicate the importance of the Lys336 side chain in binding HC to both the full-length dimer and D2^DHA^ monomer models.

### TC and DC adopt different conformations in FlgE compared to HC

Given their similar structures, it is surprising that HC appears to act as a covalent inhibitor whereas TC and DC do not. Compared to the HC pKa_1_ of ∼5, TC and DC have pKa_1_’s of 7.9 and 7.6, respectively.^41,42^ Thus, if the HC phenolate participates in 1,4 oxa-Michael addition with DHA, this species may be less abundant in TC and DC. In addition, when TC and DC are docked to the D2 DHA monomer and FlgE_FL_ dimer models (Figure S14) the top docking poses of TC (docking score = -3.893) and DC (docking score = -4.678) to the D2^DHA^ model position the phenol group away for the DHA residue. Similarly, for TC docked to the FlgE_FL_ dimer model (docking score = -4.317), the phenolate oxygen and bridging oxygen group interact with Lys336 via two 3.2 Å and 2.8 Å hydrogen bonds, but in a configuration that restrains the phenol group 6.5 Å away from DHA. The top pose of DC to the FlgE_FL_ dimer model (docking score = - 4.554) positioned the phenolate oxygen within 3.4 Å of the DHA Cβ with a 2.9 Å hydrogen bond with the ε-nitrogen of K336. These findings indicate that DC may be positioned favorably to covalently modify FlgE at higher pHs; however, this is not observed (Figure S15).

## Conclusions

Herein, we describe the development and application of a NanoLuc-based Td FlgE Lal crosslinking high-throughput screen. The assay utilized the differential reactivity of the DHA form of Td FlgE to selectively form Lal-crosslinked D1D2:D2 heterodimers without the need of cysteine-activating reagents NEM or DTNB or high ammonium sulfate concentrations. Targeting the DHA intermediate favored the discovery of covalent inhibitors reactive toward an unusual electrophilic moiety of the FlgE crosslinking reaction, thereby increasing the potential for specificity. The assay performs well on the 30 μL, 384-well scale and produced an average Z’ of 0.76 ± 0.06 across all compound plates screened in this study. Together with a false-positive counter screen and a complimentary SDS-PAGE crosslinking assay, three structurally related Lal crosslinking inhibitors, HC, TC and DC with IC_50_ values of 9.0, 47.6 and 309 μM, respectively, were discovered. HC and TC inhibit Lal crosslinking in species from three out of the four major orders of the phylum, albeit at varying potencies, which highlights the conserved nature of the Lal crosslinking mechanism but also bodes well for developing species-specific Lal crosslinking inhibitors. Biochemical data suggests that HC covalently modifies FlgE, presumably at the DHA active site, whereas TC and DC interact non-covalently with FlgE. Docking poses of each compound indicated that interaction between the phenol group and Lys336 is a conserved determinant of inhibition. This observation was supported by site-directed mutagenesis studies that revealed resistance to inhibition by HC in the Td FlgE_FL_ K336A variant compared to WT. Importantly, HC co-cultured *T. denticola* cells exhibit a dose-dependent change in FlgE oligomerization and a reduction in motility. Ultimately, because motility is a widely recognized virulence factor for pathogenic spirochetes, the results presented here establish Lal crosslinking as a legitimate target for anti-microbial therapeutic development.

## Supporting Information

Experimental methods, supporting references, construct design (Figure S1), raw chemical screening data (Figures S2, S3), SDS-PAGE crosslinking data (Figures S4 and S5), data fitting for inhibitor binding constants (Figure S6), HC and TC binding data (Figure S7), SAR results (Figure S8), DC binding data (Figure S9), FlgE crosslinking activator characterization (Figure S10), inhibitor effects on *T. denticola* cultures (Figure S11), mechanism of HC inhibition (Figure S12), docking studies of HC, TC, and DC (Figures S13,S14) and pH-dependence of HC,TC and DC inhibition (S15).

## Author Contributions

M.J.L and B.R.C. conceived the study. M.J.L. developed the assay. M.J.L. and M.D. tested compounds from the NIHCC library. M.J.L. performed the *in vitro* experiments and data analysis. K.K. grew Td cells, ran western blots and swim plate assays and analyzed *in vivo* experimental data. M.J.L. prepared figures. M.J.L. and B.R.C. wrote the manuscript, M.J.L., B.R.C., K.K. and C.L. edited the manuscript. N.W.C. provided general guidance, B.R.C. and C.L. supervised the study.

## Acknowledgements

This work was supported by NIH grants: 5R01AI148844 to B.R.C. and AI078958 and DE23080 to C.L.. The authors would also like to acknowledge the Bay Area Lyme Foundation for their support (https://www.bayarealyme.org/). The authors would like to thank and dedicate this manuscript to Dr. Faina Ryvkin (Emmanuel College, Boston MA) for guidance with *in silico* docking of the inhibitors identified in this study.

