## Supporting Information for "Inhibitors of lysinoalanine crosslinking in the flagella hook as antimicrobials against spirochetes"

[1]Department of Chemistry and Chemical Biology, Cornell University, Ithaca, NY, USA. [2]Weill Institute of Cell and Molecular Biology, Cornell University, Ithaca, NY, USA. [3]Philips Institute for Oral Health Research, Virginia Commonwealth University School of Dentistry, Richmond, VA, USA. [4]Department of Microbiology, Immunology, and Cell Biology, Robert C. Byrd Health Sciences Center, West Virginia University, Morgantown, WV, USA.

### Methods

- *Cloning, expression and protein purification*

The full-length NanoLuc plasmid was purchased from Promega. To generate the SmBit-D1D2 and LgBit-D2 fusions, *Treponema denticola* FlgE domains D1D2 (Asn91-Ser423) and D2 (Ala168-Thr344) were amplified via PCR from genomic DNA and inserted into a pET28a+ overexpression vector via Gibson cloning and confirmed by sequencing. NanoLuc LgBiT and SmBiT segments were amplified by PCR and inserted in-frame on the 3' end of the TdFlgE D2 and D1D2 expression vectors via Gibson cloning, respectively. NanoLuc SmBit-D1D2 and LgBit-D2 fusion plasmids were confirmed via DNA sequencing. See Figure S1 for the vector map and complete open reading frame amino acid sequence.

To express the NanoLuc SmBit-D1D2 and LgBit-D2 split fusions, ~200 ng of plasmid DNA was transformed into BL21(DE3) cells, plated on agar + 50 ug/mL kanamycin plates and incubated overnight at 37 ºC. Single colonies were then picked and grown in Luria broth (LB) miller media to stationary phase for 12 hours at 37 ºC with agitation and then used to inoculate 8 liters of terrific broth (TB) media to an $OD_{600}$ of 0.01-0.05. Cultures were grown at 37ºC to an $OD_{600}$ ~0.3-0.5 and then the temperature was lowered to ~22-25ºC. Protein expression was initiated via the addition of 0.1 mM isopropyl β- d-1-thiogalactopyranoside (IPTG) once the cultures reached an $OD_{600}$ of 0.6-0.8 and incubated with agitation for 16-18 hours. Cells were harvested and frozen at -20ºC.

Purification of the NanoLuc SmBit-D1D2 and LgBit-D2 split fusions proceeded via cell lysis by sonication. Cells were thawed, resuspended in 100 mL of lysis buffer (50 mM Tris pH 8, 500 mM NaCl, 5 mM imidazole) supplemented with 1 mM phenylmethanesulfonyl fluoride (PMSF) and sonicated for 12 minutes (2 sec on, 2 sec off) at 70% power. Cell debris were removed via centrifugation in a JLA30.50 rotor at 25,000 rpm for 1 hour at 4 ºC. Clarified lysate was passed through a 5 mL Nickel-NTA column pre-equilibrated with lysis buffer at a flow rate of 2-3 mL per min at 4 ºC. Bound proteins were washed with 10 column volumes of wash buffer (50 mM Tris pH 8, 500 mM NaCl, 25 mM imidazole) and eluted with elution buffer (50 mM Tris pH 8, 500 mM NaCl, 250 mM imidazole) in 1 mL fractions. Fractions containing protein were determined using Bradford reagent and pooled and concentrated down to < 5 mL using a 10 kDa MWCO spin filter. The concentrated samples were then injected into an S75 size-exclusion chromatography column (SEC, GE-26/60) pre-equilibrated with 20 mM Tris pH 7.5, 150 mM NaCl at a flow rate of 2.5 mL per min. The purity of the NanoLuc SmBit-D1D2 and LgBit-D2 split fusions were assessed via SDS-PAGE. Pure protein fractions were pooled, concentrated to < 1 mL, flash frozen in 10 µL aliquots in liquid nitrogen and stored at -80ºC. WT and mutant $FlgE_{FL}$ proteins were purified as previously described.[1,2]

To generate the dehydroalanine (DHA) form of the LgBit-D2 fusion, post-SEC fractions were pooled, concentrated to < 1 mL and mixed 1:1 with 200 mM Tris pH 9, 150 mM NaCl. Excess solid 2,5-dibromohexandiamide (DBA) was then added to the protein sample and incubated at room temperature for 30 minutes, 37 ºC for 1 hour and then 4 ºC for 12-16 hours with gently agitation. The sample was then centrifuged at 14,800 rpm for 10 minutes at 4ºC, filtered through a 0.2 µm PES filter and purified again via S75 SEC as described previously. The concentrations of all proteins were measured using the BCA assay and are reported here as the average ± the standard deviation of three technical replicates. Synthesis of DBA was performed as previously described and confirmed via $^{1}$H NMR.[1,3]

- *High-throughput screening of NIHCC and data analysis*

The high-throughput screen functioned in two main steps: First, 0.5 µL of 10 mM stock compound in dimethylsulfoxide (DMSO) was added to each well of a white, 384-well plate. The plate was then centrifuged at 1000 rpm for two minutes at room temperature. To each well, 15 µL of a mixture of 313 nM SmBit-D1D2 and 3.13 µM LgBit-D2 in buffer A (100 mM Tris pH 9, 150 mM NaCl, 5% glycerol, 1 mM EDTA, 1 mg/mL bovine serum albumin (BSA), 0.2 µm filtered) was added and the plate centrifuged. The final compound concentration in these assays was approximately 322 µM and a DMSO concentration of 3.2% (v/v). Negative controls and positive controls were supplemented with DMSO only and were comprised of SmBit-D1D2 and WT LgBit-D2 or SmBit-D1D2 and DHA LgBit-D2, respectively. The plate was then sealed with tape and incubated without agitation for 24 hours at room temperature. Following incubation, 15 µL of a 1:1000 dilution of Fz substrate is prepared in buffer B (100 mM Bis-Tris pH 6.2, 150 mM NaCl, 5% (v/v) glycerol, 1 mM EDTA, 1 mg/mL bovine serum albumin (BSA), 2 mM dithiothreitol, 0.1% (v/v) Tween20, 0.2 µm filtered) and added to each well. Following substrate addition, the plate was then sealed, centrifuged at 1000 rpm at room temperature for two minutes and the luminescence measured 5 to 30 minutes after substrate addition. Luminescence measurements (all wavelengths) were made using a SpectraMax M5 multi-mode microplate reader.

Each plate of the NIHCC was performed in duplicate. Z' values for each plate were calculated according to the equation below:

$$Z' = 1 - \frac{3(\sigma_P + \sigma_N)}{|\mu_P - \mu_N|}$$

; where $(\sigma_P + \sigma_N)$ is the sum of the standard deviations of the positive and negative controls and $|\mu_P - \mu_N|$ is the absolute value of the difference in means of the positive and negative controls. The Z' values for each plate of the NIHCC ranged from 0.63-0.82. For each plate, compounds were divided into three groups: (1) compounds with luminescence signals > $5\sigma_P$, (2) < $3\sigma_P$ and (3) < $5\sigma_P$ and > $3\sigma_P$. Compounds that fell within the first two categories and were consistent across both replicates were subjected to secondary screening protocols.

- *NanoLuc inhibition screening assay*

The following components were added to three grouped wells of a 384-well plate: well one - 0.5 μL of DMSO, well two – 0.5 μL of a 10 mM candidate compound and well three – empty. The plate was then centrifuged at 1000 rpm for 2 minutes at room temperature. Then, 15 μL of a mixture of 313 nM SmBit-D1D2 and 3.13 μM LgBit-$D2^{DHA}$ in buffer A was added and the plate centrifuged, sealed and incubated at room temperature for 24 hours. Following incubation, 0.5 μL of a 10 mM candidate compound was added to well three and allowed to incubate for 5 minutes. To each well, 15 μL of a 1:1000 dilution of Fz substrate in buffer B was then added and the plate centrifuged, and the luminescence measured. Luminescence signal was acquired at all wavelengths with a collection time of 100 ms per well. Compounds were considered hits by plotting the luminescence ratios (x-axis: $RLU^{0hr}/RLU^{DMSO}$ ratio, y-axis: $RLU^{24hr}/RLU^{DMSO}$, where well one – $RLU^{DMSO}$, well two – $RLU^{24hr}$ and well three – $RLU^{0hr}$). Inhibitors were tested further if they had $RLU^{24hr}/RLU^{DMSO}$ ratios < 0.5 and a $RLU^{0hr}/RLU^{DMSO}$ ratio < 1.2 and > 0.8. Similarly, activators were picked if they had $RLU^{24hr}/RLU^{DMSO}$ values > 2 and a $RLU^{0hr}/RLU^{DMSO}$ ratio < 1.2 and > 0.8.

- *SDS-PAGE Lal crosslinking assays*

SDS-PAGE crosslinking assays were performed as previously described with minor modifications[1]. For Coomassie-blue stained gels, 20 μM FlgE was prepared in crosslinking buffer

(50 mM Tris pH 8.5, 150 mM sodium chloride, 1 M ammonium sulfate) and incubated at 4 °C for 24 - 48 hours. Samples were then mixed 1:1 with 2x Laemmli buffer supplemented with 10% (v/v) β-mercaptoethanol (BME), heated at 95 °C for 5 minutes and electrophoreses on a 4-20% gradient tris-glycine gel. Gels were then stained with Coomassie-blue for one hour and destained to visualize protein bands. For silver-stained gels, FlgE concentrations were reduced to 2 μM and the gel stained and developed according to manufacturer's protocols. All compounds were dissolved in DMSO and 0.5-1 μL added to each crosslinking sample.

- *$IC_{50}$ determination and data fitting*

Hexachlorophene, triclosan and dichlorophen stocks were prepared in DMSO and serially diluted from 500 mM to 0.24 mM. Td FlgE samples were prepared in crosslinking buffer to a final concentration of 2 μM and 1 μL of stock compound added to yield a final concentration of 16.7 mM – 8 μM and DMSO of 3.3% (v/v). Samples were incubated at 4 °C for 48 hours, mixed 1:1 with 2x Laemmli buffer with 10% (v/v) βME and 0.5 μg of each sample resolved via SDS-PAGE. All gels were silver-stained and performed in triplicate.

Analysis of $IC_{50}$ titration data was performed using ImageJ and GraphPad Prism software.[4] Briefly, gel images were imported into FIJI in a TIF format and the band intensities of the Lal-crosslinked FlgE dimer, trimer and tetramer were measured using the gel analyzer plugin.[5] For each well, the band areas were summed and normalized against the cumulative band area of wild-type (WT) FlgE treated with DMSO for 48 hours. Normalized (norm.) cumulative band area versus compound concentration data was plotted and fit to the four-parameter [Inhibitor] vs. response equation below:

$$y = Bottom + \frac{(Top - Bottom)}{1 + \left(\frac{IC_{50}}{x}\right)^{H}}$$

; where y is the normalized, cumulative band area, bottom and top are the plateaus in the units of the y axis, x are the concentrations of compound in μM and H is the Hillslope. For hexachlorophene, the top value was constrained to one. Confidence intervals for the relative $IC_{50}$ values were computed with the likelihood ratio asymmetric method and reported as the error bars in Figure 3F (hexachlorophene [HC] and triclosan [TC]) or Figure S9 (dichlorophen [DC]).

- *Structure-activity relationship (SAR) assays*

SAR compounds were tested with the SDS-PAGE crosslinking assays. Briefly, 40 mM stocks of each compound were prepared in DMSO and mixed with 2 μM full-length Td FlgE ($FlgE_{FL}$) in 50 mM Tris pH 8.5, 150 mM sodium chloride and 1M ammonium sulfate. Samples were incubated at 4 °C for 48 hours and resolved on a 4-20% Tris-glycine SDS-PAGE gel. Protein bands were visualized by silver stain according to manufacturer's guidelines and compared to a DMSO-only carrier control.

- *Dialysis assays*

To test if HC and TC were covalent or non-covalent inhibitors, two dialysis experiments were performed – one using full-length Td FlgE (Figure S7) and the other with truncated Td FlgE D1D2 and D2 domains (Figure 4). For $FlgE_{FL}$, 20 μM (1 mg/mL) protein was prepared in crosslinking buffer and mixed with 1 mM HC, 1 mM TC or 100 mM βME. Samples were incubated at 4 °C for 48 hours to promote Lal crosslinking and then dialyzed against crosslinking buffer without compound at 4 °C for 48 hours. After dialysis, samples were dialyzed against 150 mM sodium phosphate pH 6.5 and the concentrations measured via BCA assay. Lal crosslinking for the 0-, 2- and 4-day timepoints were monitored via SDS-PAGE and visualized via silver-stain. For each sample, approximately 1 μg of protein was loaded per well and each gel was performed in triplicate. Lal crosslinked oligomers were quantified using ImageJ as described previously.[1] This assay was also performed with truncated Td FlgE D1D2 and $D2^{DHA}$ domains with some modifications. Briefly, 100 μM $D2^{DHA}$ was prepared in 20 mM Tris pH 9, 150 mM sodium chloride and supplemented with 1 mM HC, 1 mM TC or 100 mM βME. Samples were incubated for 24 hours at 4 °C and then dialyzed against 20 mM Tris pH 7.5, 150 mM sodium chloride. Protein concentrations were measured using the BCA assay and then mixed in a 4:1 ratio with Td FlgE D1D2 (100 μM $D2^{DHA}$: 25 μM D1D2) in 100 mM Tris pH 9, 150 mM sodium chloride and incubated at 4 °C for 24 hours. Lal crosslinking for the 0- and 1-day timepoints were monitored via SDS-PAGE and visualized via Coomassie-blue staining. For each sample, approximately 10 μg of total protein was loaded per well and each gel was performed in triplicate. Lal-crosslinked dimers were quantified using ImageJ as described previously.[1] The pH-dependence of HC, TC and DC inhibition, 100 μM $D2^{DHA}$ was prepared as described above, except at pH 7, 8 and 9. Similarly, the time-dependence of WT $FlgE_{FL}$ inhibition was determined via pre-treatment of 20 μM $FlgE_{FL}$ with 500 μM HC for varying time points (0 min, 10 min, 30 min). Samples were then dialyzed, crosslinked and analyzed as described.

- *T. denticola cell culture and FlgE western blot analysis*

All TYGVS media was prepared fresh and incubated anaerobically for at least 24 hours prior to use. Stocks of wild-type *T. denticola* ATCC 34505 cells in TYGVS media supplemented with 20% glycerol (v/v) were thawed and inoculated into 3 mL of media at a density of $10^5$ cells per mL. To each stock, 5 μL of hexachlorophene was added to yield a final concentration of 83.3 μM. For DMSO only control samples, 5 μL of DMSO was added in lieu of HC. Cells were grown at 37 °C anaerobically and passaged for four generations, then 150 *T. denticola* cells were plated on 75% TYGVS:25% (v/v) phosphate-buffered saline (50 mM sodium phosphate pH 7.5, 150 mM sodium chloride) plates prepared with 0.35% (w/v) sea plaque agarose supplemented with DMSO or 83.3 μM HC. Plates were incubated anaerobically at 37 °C for seven days and individual colonies picked and grown in TYGVS media with DMSO (2 colonies, D1-D2) or 83.3 μM HC (5 colonies, H1-H5) to stationary phase and analyzed by western blot.

Cells were lysed via sonication and protein concentration determined via BCA assay. Samples were normalized to total protein concentration, mixed 1:1 with 2x Laemmli buffer supplemented with excess BME and heated at 95 °C. Samples were then centrifuged and loaded on an SDS-PAGE gel and electrophoresed at 200V in Tris-glycine SDS running buffer. The gels were rinsed with transfer buffer (25 mM Tris, 192 mM glycine, pH 8.3) and transferred to a 0.2 um PVDF membrane. Membranes were then blocked in 5% (w/v) skim milk in tris-buffered saline supplemented with 0.2% (v/v) Tween20 (TBST) for one hour at room temperature and incubated with a 1:1000 (v/v) dilution of FlgE antibody (αFlgE) that was raised in rats in 5% (w/v) skim milk/TBST for one hour at room temperature. Membranes were then washed three times with TBST, incubated with 1:5000 (v/v) dilution of αRat HRP-IgG in 5% (w/v) skim milk/TBST for one hour and washed three times with TBST. Chemiluminescent bands were visualized via the addition of HRP chemiluminescent substrate and imaged within 30 seconds.

- *T. denticola swimming plate assays*

Swimming plate assays were performed as previously described with minor modifications.[6] Plates containing 50% (v/v) TYGVS:50% (v/v) PBS with 0.35% (w/v) sea plaque agarose were prepared and incubated in an anerobic chamber for 24 hours. Wild-type, ΔTap1(a non-motile *fliK* deletion mutant) and HC-treated (colonies 1 and 2, 83.3 μM) *T. denticola* cell were grown anaerobically at 37°C until the cell density reaches ~5x$10^8$ – $10^9$ cells per mL. Cells were then centrifuged and resuspended in ~20 μL of TYGVS media normalizing to cell density. Each plate was inoculated with 4 μL, dried for 15-30 minutes in a biosafety-hood and transferred to a 37°C anerobic incubator for three days. *T. denticola* swimming ring diameters were measured and recorded as the average ± standard deviation of four replicates.

- <u>*FlgE monomer and dimer model generation and compound docking*</u>

To generate a mixed Cys/DHA dimer model for docking analysis, a wild-type model of Td FlgE was generated using AlphaFold3.[7] The D2 domain (Ala168-Thr334) of one FlgE monomer was replaced with the DHA D2 domain crystal structure determined previously (PDB: 6NDT) to generate a mixed Cys/DHA dimer model.[1] To generate the D2 DHA monomer model, the D2 DHA crystal structure was used without modification.[1] First, the models were prepared using protein preparation wizard.[8] A pH of 8.5 was used based on the experimental conditions of our *in vitro* SDS-PAGE crosslinking assays. Prepared model energy minimization was performed using Macromodel[9] minimization using the following parameters: OPLS4[10] force-field, implicit solvation with a constant dielectric of 78, charges were calculated from the force-field with normal cutoffs, PRCG minimization method, 100,000 maximum iterations and models were allowed to converge on a gradient with a threshold of 0.001. Hexachlorophene, triclosan and dichlorophen were imported into Maestro[11] and prepared using LigPrep[11] at a pH of 8.5. Docking grids (12x12x12 $Å^3$) were generated in Glide[12] centered on residue DHA-178. Docking poses were generated using Glide Dock using standard-precision docking using default parameters with minor modifications. Ligand-protein hydrogen bonds were rewarded, and halogens were allowed to participate as hydrogen bond acceptors. At least 2000 poses were considered per ligand and all poses were subject to post-docking energy minimization using an OPLS4 force field.[10] The top pose of each ligand was output for analysis.

**Supplemental Figures**

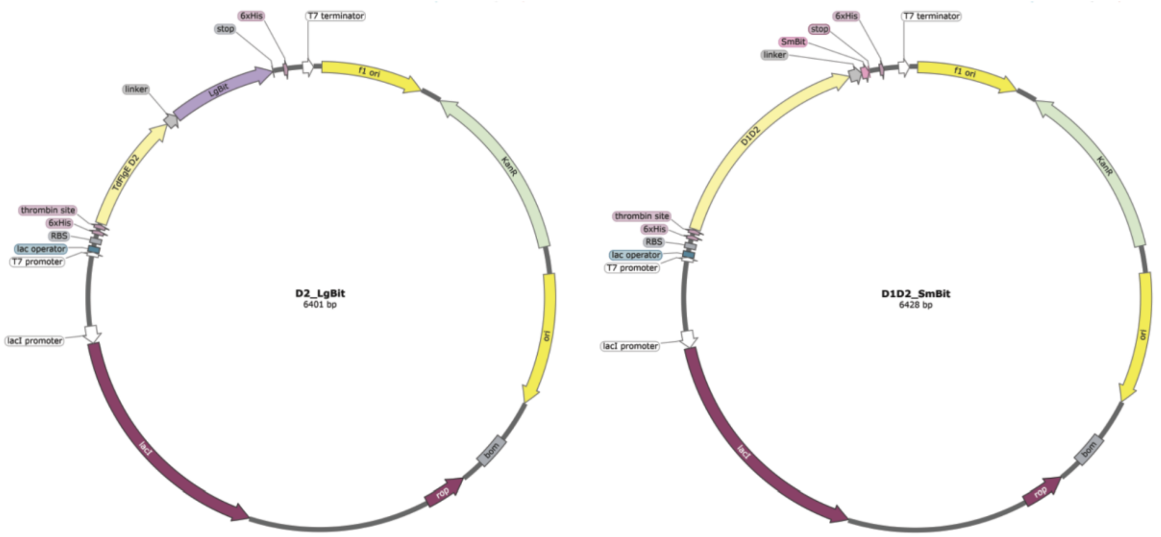


D2_LgBit sequence:

MGSSHHHHHHSSGLVPRGSHMMAKATTSVNYACNLDKRLPELPEGANRAQILESTWSTEFKVYDSFGEAHELQIDFARVPGEVNAWRAT
VNVDPTNADATATRVGIGTTDGVQNSFIVRFDNNGHLASVTDTAGNVTSPAGQVLVQISYNVVGANPDEAGAPTRHTFDVNLGEIGTSKNT
ITQFSDKSTTKAYEQDGYTVSQGSSGGGGSGGGGSSG**VFTLEDFVGDWEQTAAYNLDQVLEQGGVSSLLQNLAVSVTPIQRIVRSGENA**
**LKIDIHVIIPYEGLSADQMAQIEEVFKVVYPVDDHHFKVILPYGTLVIDGVTPNMLNYFGRPYEGIAVFDGKKITVTGTLWNGNKIIDERLITPD**
**GSMLFRVTINS**

D1D2_SmBit sequence:

MGSSHHHHHHSSGLVPRGSHMMNTDLAIQGNGFFILKDGEKTFYTRAGAFGIDKEGTLVNPANGMRVQGWMAEEAEGFRIINTSGQTED
LNIPIGQKLDAKATTSVNYACNLDKRLPELPEGANRAQILESTWSTEFKVYDSFGEAHELQIDFARVPGEVNAWRATVNVDPTNADATATRV
GIGTTDGVQNSFIVRFDNNGHLASVTDTAGNVTSPAGQVLVQISYNVVGANPDEAGAPTRHTFDVNLGEIGTSKNTITQFSDKSTTKAYEQD
GYTLGYLENFRIDQSGIITGVYSNGVRQEIGQIAMAGFANQGGLEKAGQNTYVQSNNSGIANVSTSGTVGKGYFIGGTLEMSVSQGSSGGG
GSGGGGSSG**VTGYRLFEEIL**

Figure S1: Constructing LgBiT-D2 and SmBiT-D1D2 fusions.

Split NanoLuc domains are fused to the C-termini of the Td FlgE D2 and D1D2 domains and denoted with bold and underlined text.

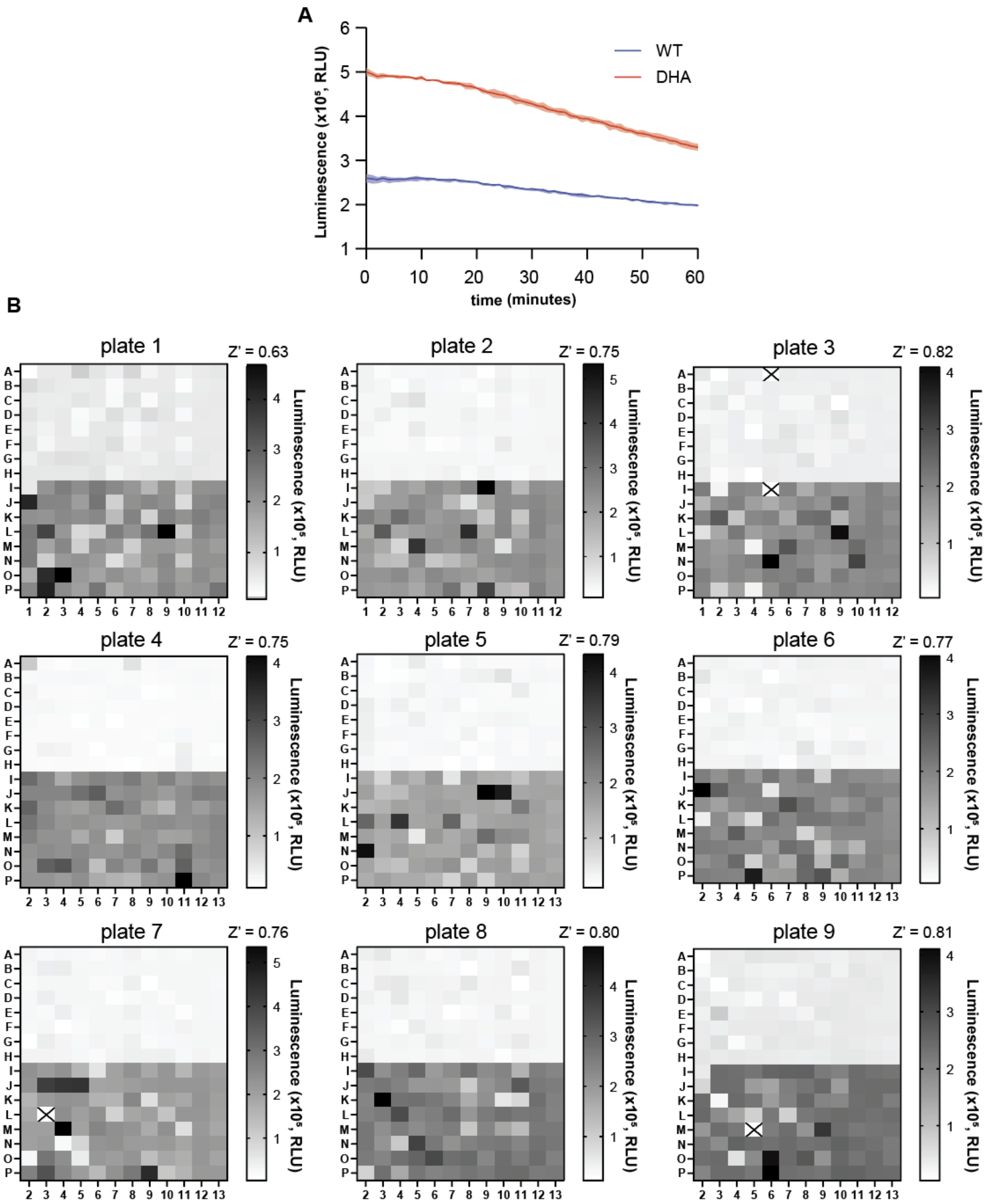
A
Luminescence (x10⁵, RLU)
time (minutes)
WT
DHA
B
plate 1
Z' = 0.63
plate 2
Z' = 0.75
plate 3
Z' = 0.82
plate 4
Z' = 0.75
plate 5
Z' = 0.79
plate 6
Z' = 0.77
plate 7
Z' = 0.76
plate 8
Z' = 0.80
plate 9
Z' = 0.81
Luminescence (x10⁵, RLU)

Figure S2: Raw chemiluminescence data from NanoLuc-FlgE high-throughput screen.

**A)** Kinetic trace of luminescence signal versus time of negative (WT, blue) and positive (DHA, red) controls. For each sample, the signal was measured and reported as the average signal ± the standard deviation of three technical replicates. **B)** Raw luminescent data from screening the National Institutes of Health Clinical Compound (NIHCC) library. Each plate is divided into two segments with LgBiT-D2$^{WT}$/SmBiT-D1D2 + compound in rows A-H and LgBiT-D2$^{DHA}$/SmBiT-D1D2 + compound in rows I-P. Z' values for each plate are calculated from negative and positive controls in rows A23/24-H23/24 and I23/24-P23/24, respectively (n=16 per control per plate). Outlier wells are marked by a black X.

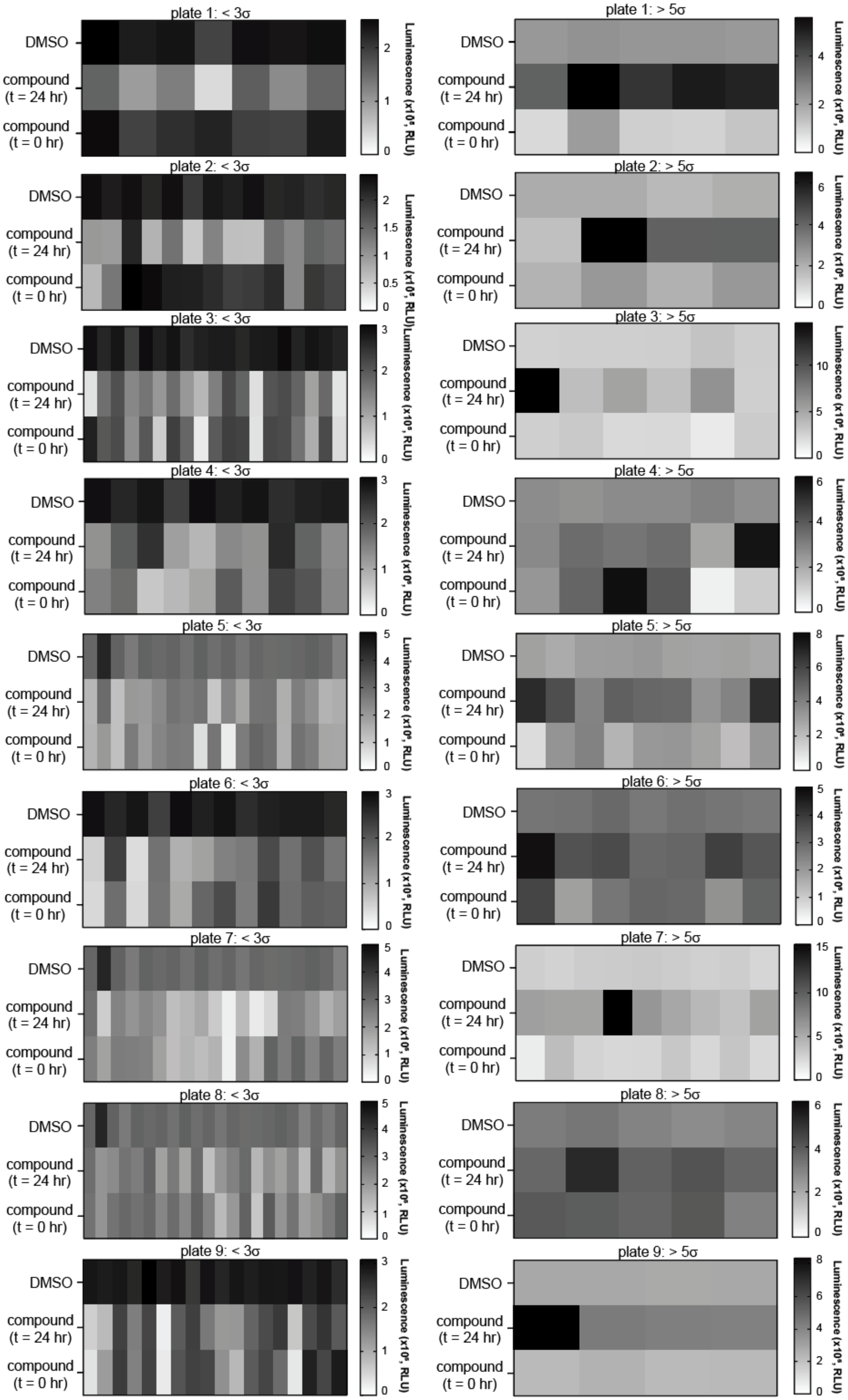
plate 1: < 3σ
plate 1: > 5σ
plate 2: < 3σ
plate 2: > 5σ
plate 3: < 3σ
plate 3: > 5σ
plate 4: < 3σ
plate 4: > 5σ
plate 5: < 3σ
plate 5: > 5σ
plate 6: < 3σ
plate 6: > 5σ
plate 7: < 3σ
plate 7: > 5σ
plate 8: < 3σ
plate 8: > 5σ
plate 9: < 3σ
plate 9: > 5σ
DMSO
compound (t = 24 hr)
compound (t = 0 hr)

Figure S3: NanoLuc inhibition screening assay

Raw luminescent data from our secondary screening procedure to remove false-positive hits arising from NanoLuc inhibition. Compounds with reproducible (n = 2) increased (>5σ) or decreased (<3σ) luminescent signals identified by the NanoLuc high-throughput screen were collected and rescreened. For each plate, a mixture of LgBiT-D2$^{DHA}$ and SmBiT-D1D2 was incubated with DMSO (top row), compound for 24 hours (middle), or compound for 0 hours (bottom). The ratio of the bottom and top rows (control/control) and the middle and top rows (treated/control) were plotted as the x and y coordinates, respectively, in Figure 2C-D.

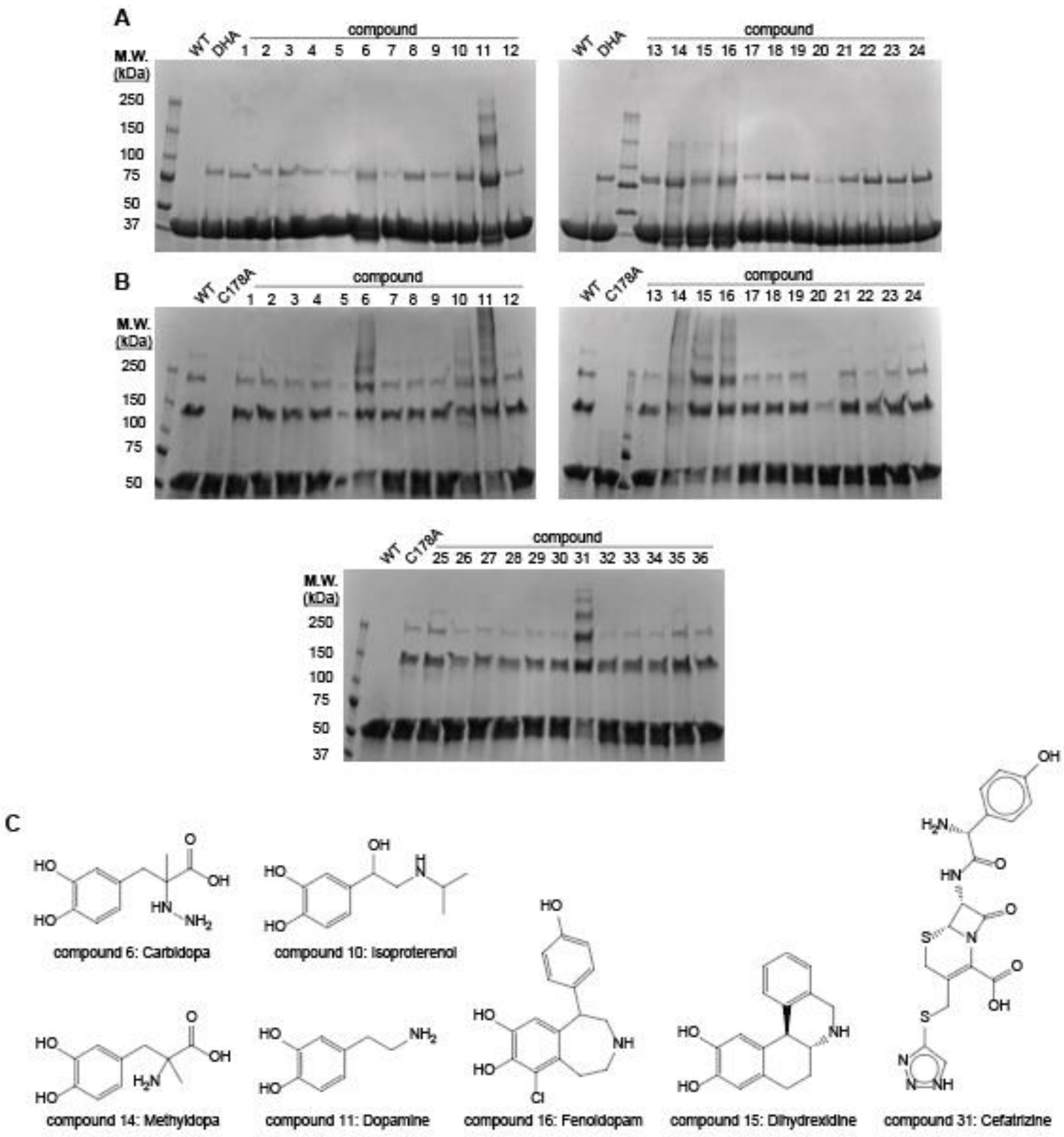


Figure S4: *in vitro* SDS-PAGE crosslinking assay on verified NIHCC compound hits

**A)** LgBiT-D2$^{DHA}$ and SmBiT-D1D2 incubated with candidate compounds following NanoLuc inhibition screening. For each sample, 10 μM LgBiT-D2$^{DHA}$ and 50 μM SmBiT-D1D2 were incubated with 322 μM compound for 24 hours at room temperature. A WT and DHA sample consisting of 50 μM SmBiT-D1D2 and 10 μM WT or DHA LgBiT-D2 and 3.2% (v/v) DMSO were included as negative and positive controls, respectively. **B)** Screening of the same compounds in (A) except against 20 μM Td $FlgE_{FL}$. WT and C178A Td $FlgE_{FL}$ were included in these assays as a positive and negative control, respectively. Gels in (A) and (B) were stained with Coomassie-blue. **C)** Identity and structure of compounds discovered in the assays that activate Td FlgE (either truncated or full length).

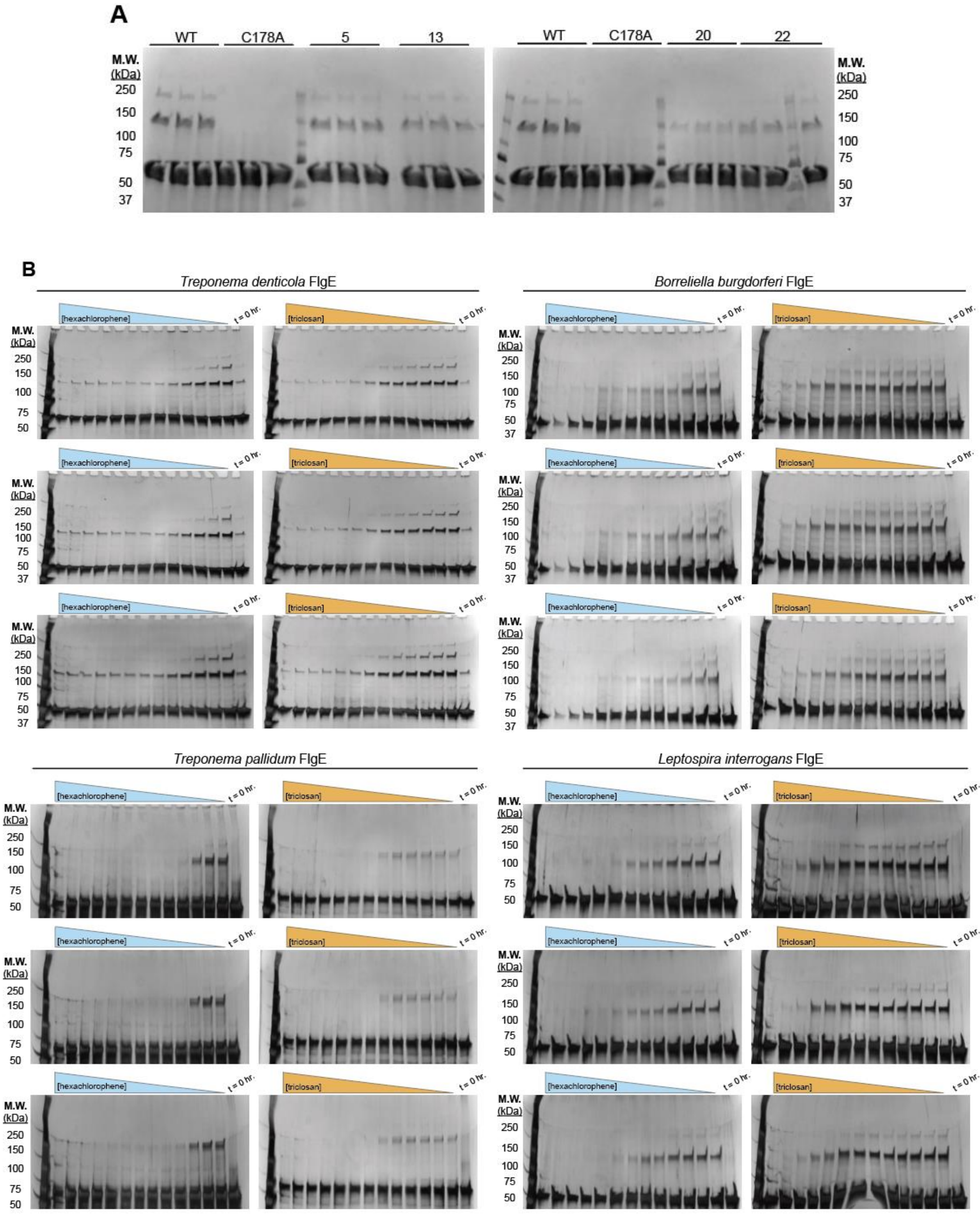

A
WT
C178A
5
13
WT
C178A
20
22
M.W. (kDa)
250
150
100
75
50
37
B
Treponema denticola FlgE
Borreliella burgdorferi FlgE
Treponema pallidum FlgE
Leptospira interrogans FlgE
[hexachlorophene]
[triclosan]
t = 0 hr.

Figure S5: SDS-PAGE crosslinking assay gels with $FlgE_{FL}$

**A)** Replicates of top NIHCC inhibitor candidates. For each sample, 20 μM $FlgE_{FL}$ was incubated with 333 μM compound for 48 hours at 4°C. Inhibition was measured by quantification of Coomassie-stained Lal crosslinked high-molecular weight complex (HMWC) bands for each sample and compared to the wild-type sample (DMSO only). **B)** $IC_{50}$ replicate gels for HC and TC titration experiments. For each sample, 2 μM Td, Tp, Bb and Li $FlgE_{FL}$ was incubated for 48 hours (96 hours for Bb) with varying concentrations of Hex and Tri. Silver-stained Lal cross-linked HMW bands were measured and compared to t = 0-hour sample. For each concentration, the cumulative HMWC band area was normalized against a DMSO only sample and reported as the average normalized cumulative Lal crosslink band area ± the standard deviation of three technical replicates. $IC_{50}$ curves are reported in Figure 3D-E and values shown in Figure 3F.

**A**

$$y = Bottom + \frac{(Top - Bottom)}{1 + \left(\frac{IC_{50}}{x}\right)^{H}}$$

**B**

Hexachlorophene data fitting statistics:

| | Nonlin fit<br>Table of results | A<br>Td | B<br>Bb | C<br>Tp | D<br>Li |
|---|---|---|---|---|---|
| 1 | **[Inhibitor] vs. response -- Variable slope (four parameters)** | | | | |
| 2 | **Best-fit values** | | | | |
| 3 | Bottom | 0.1675 | 0.07873 | 0.05695 | 0.003923 |
| 4 | Top | = 1.000 | = 1.000 | = 1.000 | = 1.000 |
| 5 | IC50 | 8.999 | 14.19 | 9.374 | 48.33 |
| 6 | HillSlope | -1.427 | -0.9901 | -3.072 | -1.143 |
| 7 | logIC50 | 0.9542 | 1.152 | 0.9719 | 1.684 |
| 8 | Span | 0.8325 | 0.9213 | 0.9431 | 0.9961 |
| 9 | **95% CI (profile likelihood)** | | | | |
| 10 | Bottom | 0.1454 to 0.1891 | 0.01822 to 0.1296 | 0.02813 to 0.08540 | -0.04004 to 0.04452 |
| 11 | IC50 | 7.870 to 10.25 | 10.69 to 19.26 | 8.442 to 10.45 | 40.18 to 58.60 |
| 12 | HillSlope | -1.710 to -1.191 | -1.390 to -0.7146 | -4.396 to -2.274 | -1.359 to -0.9672 |
| 13 | logIC50 | 0.8960 to 1.011 | 1.029 to 1.285 | 0.9264 to 1.019 | 1.604 to 1.768 |
| 14 | **Goodness of Fit** | | | | |
| 15 | Degrees of Freedom | 36 | 36 | 36 | 36 |
| 16 | R squared | 0.9712 | 0.9283 | 0.9626 | 0.9751 |
| 17 | Sum of Squares | 0.08352 | 0.2776 | 0.1784 | 0.1499 |
| 18 | Sy.x | 0.04817 | 0.08781 | 0.07040 | 0.06453 |
| 19 | **Constraints** | | | | |
| 20 | Top | Top = 1 | Top = 1 | Top = 1 | Top = 1 |
| 21 | IC50 | IC50 > 0 | IC50 > 0 | IC50 > 0 | IC50 > 0 |
| 22 | | | | | |
| 23 | **Number of points** | | | | |
| 24 | # of X values | 39 | 39 | 39 | 39 |
| 25 | # Y values analyzed | 39 | 39 | 39 | 39 |
| 26 | | | | | |

Triclosan data fitting statistics:

| | Nonlin fit<br>Table of results | A<br>Td | B<br>Bb | C<br>Tp | D<br>Li |
|---|---|---|---|---|---|
| 1 | **[Inhibitor] vs. response -- Variable slope (four parameters)** | | | | |
| 2 | **Best-fit values** | | | | |
| 3 | Bottom | 0.08184 | -0.1053 | 0.01475 | -0.02921 |
| 4 | Top | 0.8987 | 1.101 | 0.8584 | 0.9148 |
| 5 | IC50 | 61.92 | 1459 | 64.66 | 909.3 |
| 6 | HillSlope | -1.227 | -1.111 | -4.511 | -1.262 |
| 7 | logIC50 | 1.792 | 3.164 | 1.811 | 2.959 |
| 8 | Span | 0.8169 | 1.207 | 0.8437 | 0.9441 |
| 9 | **95% CI (profile likelihood)** | | | | |
| 10 | Bottom | 0.001357 to 0.09804 | -0.6807 to 0.1679 | -0.02552 to 0.05203 | -0.6355 to 0.09793 |
| 11 | Top | ??? to 1.068 | 1.018 to 1.154 | 0.8359 to 0.9448 | 0.8790 to 1.013 |
| 12 | IC50 | 35.28 to 65.91 | 851.4 to 4560 | 56.92 to 72.38 | 598.5 to 3236 |
| 13 | HillSlope | -1.307 to -0.7387 | -2.297 to -0.7336 | ??? to -2.586 | -1.847 to -0.5955 |
| 14 | logIC50 | 1.548 to 1.819 | 2.930 to 3.659 | 1.755 to 1.860 | 2.777 to 3.510 |
| 15 | **Goodness of Fit** | | | | |
| 16 | Degrees of Freedom | 35 | 35 | 35 | 35 |
| 17 | R squared | 0.9824 | 0.9027 | 0.9676 | 0.9302 |
| 18 | Sum of Squares | 0.07750 | 0.5401 | 0.2170 | 0.3201 |
| 19 | Sy.x | 0.04706 | 0.1242 | 0.07875 | 0.09563 |
| 20 | **Constraints** | | | | |
| 21 | IC50 | IC50 > 0 | IC50 > 0 | IC50 > 0 | IC50 > 0 |
| 22 | | | | | |
| 23 | **Number of points** | | | | |
| 24 | # of X values | 39 | 39 | 39 | 39 |
| 25 | # Y values analyzed | 39 | 39 | 39 | 39 |

Figure S6: Data fitting parameters for hexachlorophene and triclosan titrations

**A)** Average normalized cumulative Lal-crosslinked band area data for Td, Tp, Bb and Li FlgE were fit to the four-parameter [Inhibitor] vs. response equation with GraphPad Prism. Confidence intervals were computed using the likelihood ratio asymmetric method. **B)** Fitting parameters and values for HC and TC titrations.

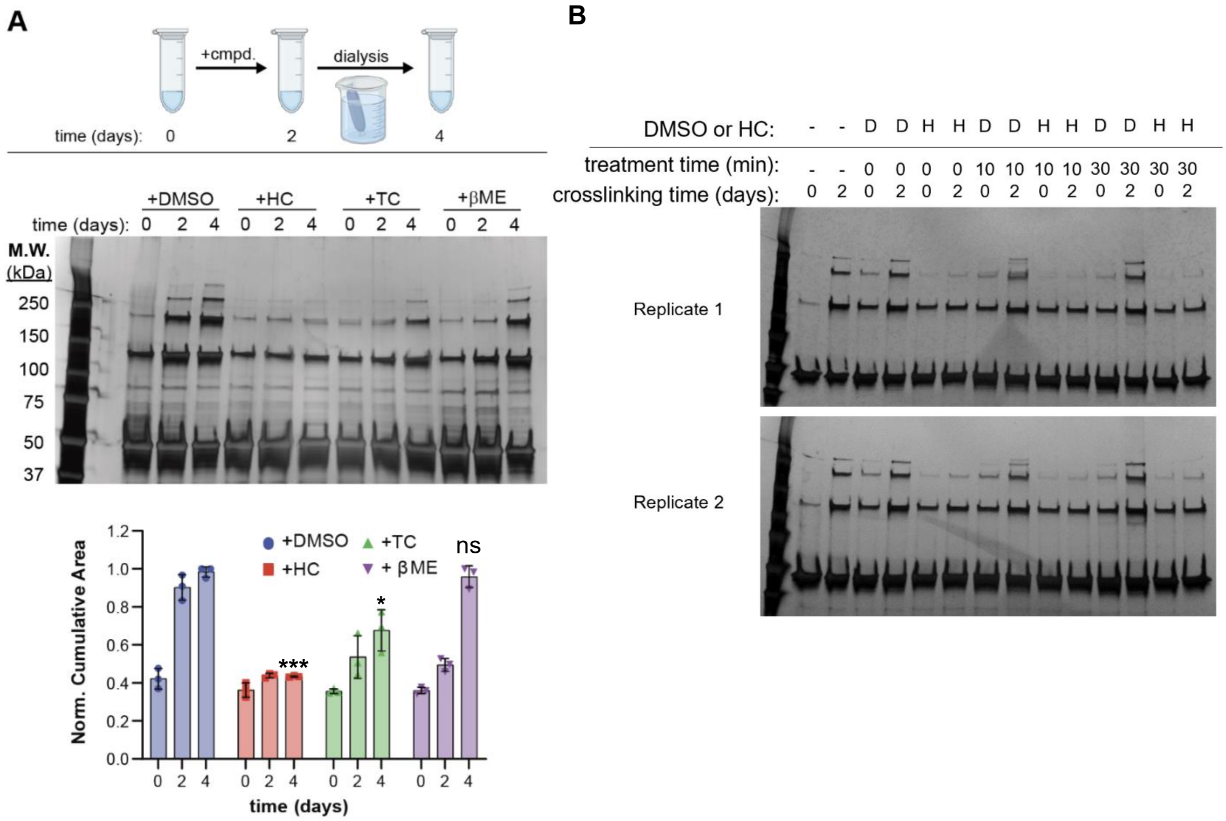


Figure S7: Differential inhibition by HC and TC

**(A)** *In vitro* SDS-PAGE LaI crosslinking assay with 2 µM $FlgE_{FL}$ incubated with 1 mM HC, TC or 100 mM βME. (top) Overview of experimental design of assay. Each sample was crosslinked in the presence of DMSO or compound for two days at 4 ºC. Samples were then dialyzed against fresh crosslinking buffer to remove DMSO, HC, TC or βME and crosslinking continued for another 48 hours. Samples were taken every two days and the presence of LaI-crosslinked FlgE monomers was monitored via silver-stain SDS-PAGE. Statistical significance was calculated using a two-tailed Welch's t-test ($p<0.05$*, 0.01** and 0.001****) and comparisons made between the day 4 time-points for DMSO and HC, TC and βME-treated samples. **(B)** HC inhibits LaI crosslinking in $FlgE_{FL}$ samples in a little as ten minutes. For each sample, 2 µM $FlgE_{FL}$ was incubated with 500 µM HC in crosslinking buffer for varying time-points and then dialyzed to remove compound for 24 hours. HC-treated samples were crosslinked for two days and LaI formation monitored via silver-stain SDS-PAGE.

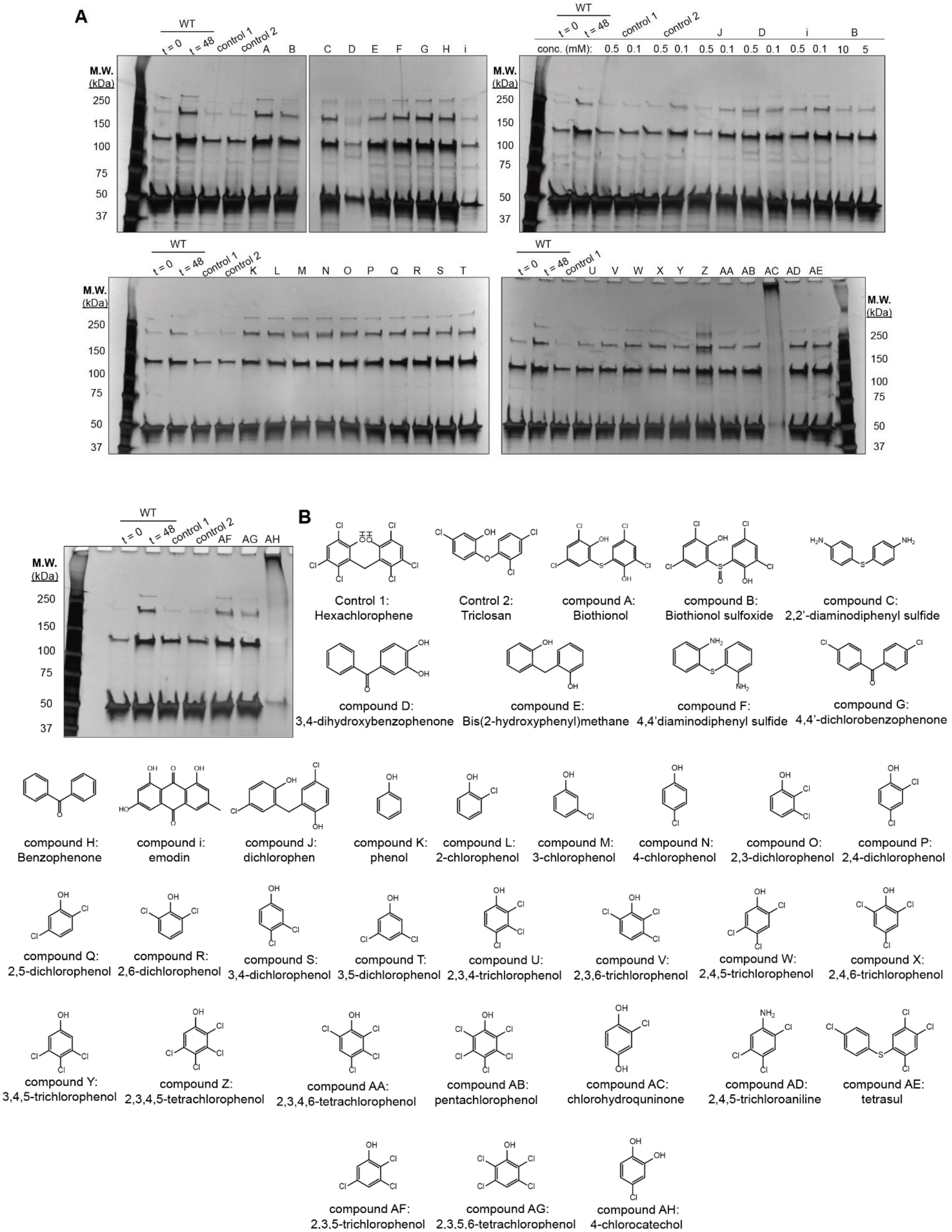
A
WT
t = 0
t = 48
control 1
control 2
A B C D E F G H i
M.W. (kDa)
250
150
100
75
50
37
conc. (mM):
J D i B
0.5 0.1 0.5 0.1 0.5 0.1 0.5 0.1 0.5 0.1 10 5
K L M N O P Q R S T
U V W X Y Z AA AB AC AD AE
AF AG AH
B
Control 1: Hexachlorophene
Control 2: Triclosan
compound A: Biothionol
compound B: Biothionol sulfoxide
compound C: 2,2'-diaminodiphenyl sulfide
compound D: 3,4-dihydroxybenzophenone
compound E: Bis(2-hydroxyphenyl)methane
compound F: 4,4'diaminodiphenyl sulfide
compound G: 4,4'-dichlorobenzophenone
compound H: Benzophenone
compound i: emodin
compound J: dichlorophen
compound K: phenol
compound L: 2-chlorophenol
compound M: 3-chlorophenol
compound N: 4-chlorophenol
compound O: 2,3-dichlorophenol
compound P: 2,4-dichlorophenol
compound Q: 2,5-dichlorophenol
compound R: 2,6-dichlorophenol
compound S: 3,4-dichlorophenol
compound T: 3,5-dichlorophenol
compound U: 2,3,4-trichlorophenol
compound V: 2,3,6-trichlorophenol
compound W: 2,4,5-trichlorophenol
compound X: 2,4,6-trichlorophenol
compound Y: 3,4,5-trichlorophenol
compound Z: 2,3,4,5-tetrachlorophenol
compound AA: 2,3,4,6-tetrachlorophenol
compound AB: pentachlorophenol
compound AC: chlorohydroquninone
compound AD: 2,4,5-trichloroaniline
compound AE: tetrasul
compound AF: 2,3,5-trichlorophenol
compound AG: 2,3,5,6-tetrachlorophenol
compound AH: 4-chlorocatechol

Figure S8: Structure-activity relationship (SAR) studies for HC and TC

**A)** *in vitro* SDS-PAGE Lal crosslinking assay gels against various compounds. For each experiment, 2 μM Td $FlgE_{FL}$ was incubated with 1 mM compound for 48 hours at 4 ºC. Lal-crosslinked HMWC bands were visualized via silver-stain and compared to 0hr and 48hr WT Td $FlgE_{FL}$ + 2.5% (v/v) DMSO samples as negative and positive controls respectively. For each assay, inhibition controls of 1mM HC (control 1) and/or TC (control 2) were included for comparison. **B)** Name and structure of compounds tested in our SAR studies.

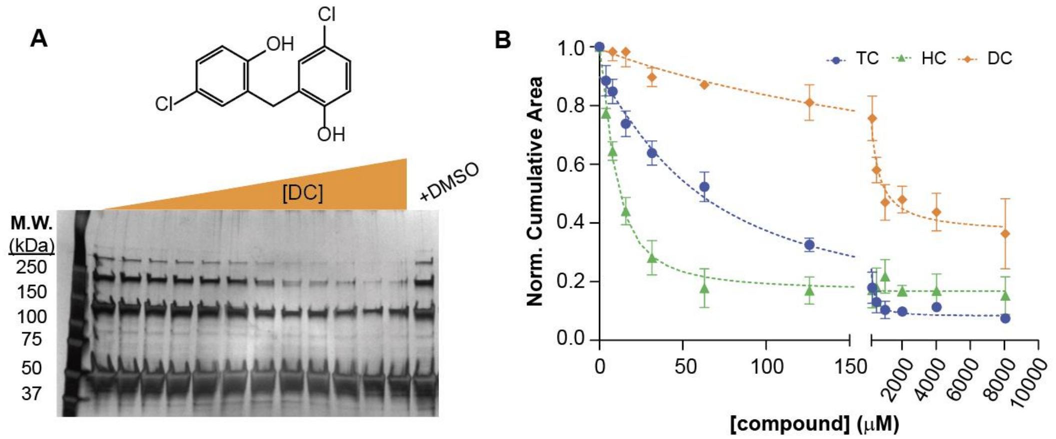


C

| | Nonlin fit<br>Table of results | A<br>dichlorophen |
|---|---|---|
| 1 | **[Inhibitor] vs. response -- Variable slope (four parameters)** | |
| 2 | **Best-fit values** | |
| 3 | Bottom | 0.3617 |
| 4 | Top | 0.9935 |
| 5 | IC50 | 308.7 |
| 6 | HillSlope | -0.9755 |
| 7 | logIC50 | 2.489 |
| 8 | Span | 0.6318 |
| 9 | **95% CI (profile likelihood)** | |
| 10 | Bottom | 0.2565 to 0.4246 |
| 11 | Top | 0.9480 to 1.056 |
| 12 | IC50 | 211.1 to 526.6 |
| 13 | HillSlope | -1.440 to -0.6614 |
| 14 | logIC50 | 2.325 to 2.721 |
| 15 | **Goodness of Fit** | |
| 16 | Degrees of Freedom | 32 |
| 17 | R squared | 0.9453 |
| 18 | Sum of Squares | 0.1074 |
| 19 | Sy.x | 0.05793 |
| 20 | **Constraints** | |
| 21 | IC50 | IC50 > 0 |
| 22 | | |
| 23 | **Number of points** | |
| 24 | # of X values | 36 |
| 25 | # Y values analyzed | 36 |

Figure S9: $IC_{50}$ determination of DC

**A)** *in vitro* SDS-PAGE Lal-crosslinking assay with WT Td $FlgE_{FL}$ in the presence of varying DC concentrations. Samples of 2 μM Td FlgE were incubated at 4ºC for 48 hours and Lal-crosslinked HMW bands visualized by silver-stain. A DMSO only sample was included as a negative (no inhibition) control. **B)** IC50 curve data for DC compared to HC and TC. Normalized cumulative band areas for each DC concentration are reported as the average ± standard deviation of three technical replicates. **C)** DC four-parameter [Inhibitor] vs. response equation data fitting parameters and values. Data was fit using GraphPad Prism as described in Figure S6.[4]

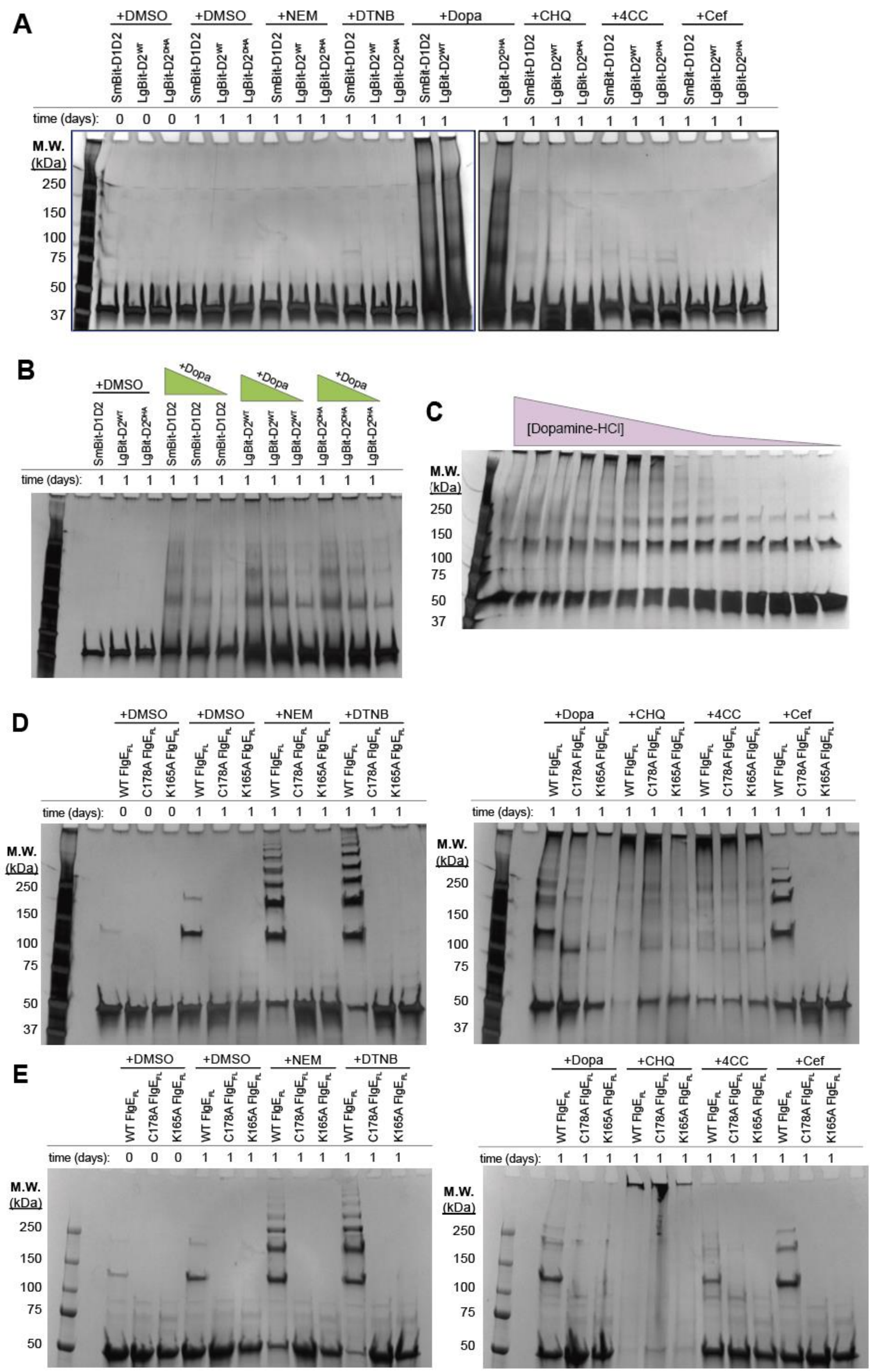
A
+DMSO
+DMSO
+NEM
+DTNB
+Dopa
+CHQ
+4CC
+Cef
SmBit-D1D2
LgBit-D2WT
LgBit-D2DHA
time (days):
M.W. (kDa)
250
150
100
75
50
37
B
+DMSO
+Dopa
C
[Dopamine-HCl]
D
WT FlgE FL
C178A FlgE FL
K165A FlgE FL
E

Figure S10: Characterization of FlgE activators

**A)** SDS-PAGE Lal crosslinking assay with DHA, WT $FlgE_{D2}$-LgBit and $FlgE_{D1D2}$-SmBit. Each sample was incubated with the following activators identified in the HTS and SAR studies: Dopa – dopamine HCl, CHQ – chlorohydroquinone, 4CC – 4-chlorocatechol and Cef – cefatrizine. N-ethylmaleimide (NEM) and 5,5′-dithiobis-[2-nitrobenzoic acid] (DTNB) were included as cysteine-reactive compound controls. For each sample, 2 μM total protein was incubated with 1 mM compound at 4 ºC for 24 hours. DMSO-only samples at 0- and 24-hour timepoints were used as controls to monitor Lal crosslink formation over the course of the experiment. **B)** Dopa titration with 2 μM LgBiT-$D2^{DHA}$, LgBiT-$D2^{WT}$ and SmBiT-D1D2. Dopa concentrations tested range from 167 – 42 2 μM. **C)** Dopa titration with 2 μM WT Td $FlgE_{FL}$. Dopamine concentrations range from 0-16 mM. **D)** SDS-PAGE Lal crosslinking assay with full-length WT, C178 and K165A Td $FlgE_{FL}$. Each sample was incubated with DMSO (3.3% (v/v), control), NEM, DTNB, Dopa, CHQ, 4CC or Cef (1 mM final concentration) for 24 hours at 4ºC. Protein bands of SDS-PAGE gels in A-D were visualized by silver stain. **E)** Identical experiments as shown in (D) except for the following differences: total FlgE concentrations used were lowered to 20 μM and protein bands were visualized by Coomassie stain.

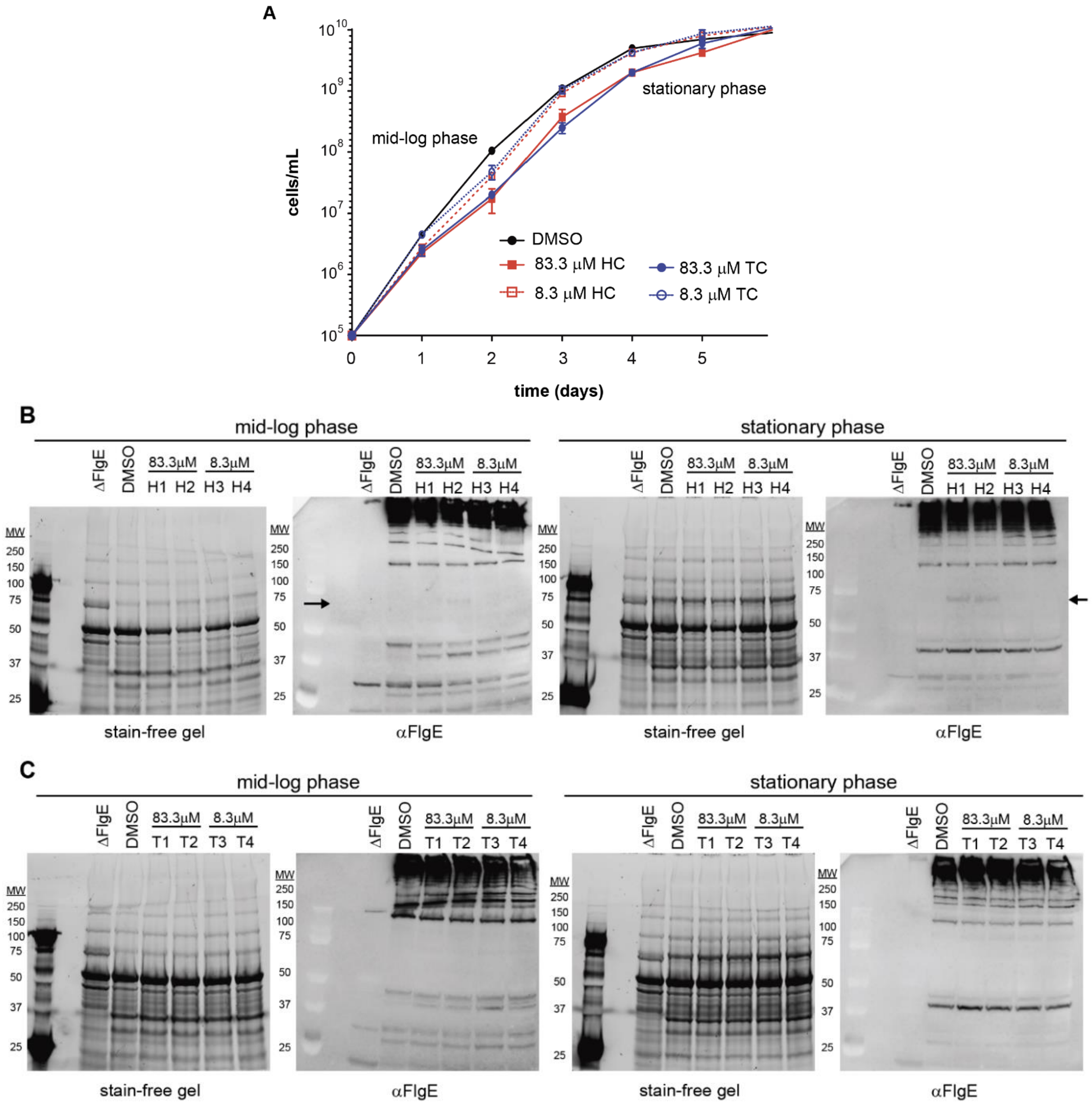


Figure S11: *T. denticola* cell cultures in the presence of HC and TC

**A)** *T. denticola* growth curves in the presence of DMSO (control, black line), HC (83.3 μM – red, solid, 8.3 μM – red, dotted) and TC (83.3 μM – blue, solid, 8.3 μM – blue, dotted). Cell densities are reported as the average ± the standard deviation of three biological replicates. **B)** Stain-free SDS-PAGE gels (left) and western blots probed against αFlgE (right) of whole-cell lysate of Td cells co-cultured with DMSO, 83.3 μM, or 8.3 μM HC. The ΔFlgE strain was cultured without DMSO as a negative (no crosslinking/FlgE) control. Td cell lysate was normalized to cell densities (cells/mL) and repeated in duplicate. Cells were analyzed as two different time points: (1) at mid-log phase (left) and (2) stationary phase (right). Differential αFlgE-reactive bands are denoted by a black arrow. **C)** stain-free SDS-PAGE gels (left) and αFlgE western blots (right) of whole-cell lysate of Td cells co-cultured with DMSO, 83.3 μM, or 8.3 μM TC. Experiment design identical to (B).

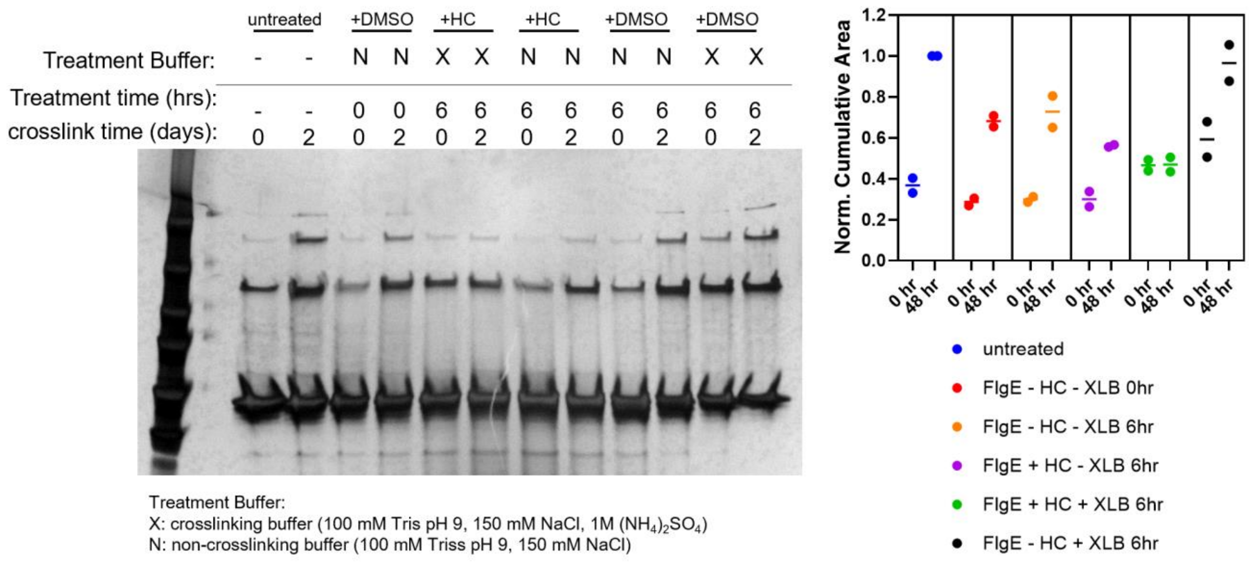


<u>Figure S12: HC requires active Lal crosslinking conditions to inhibit FlgE</u>

(left) Silver-stained SDS-PAGE gel of 2 μM Td FlgE$_{FL}$ pre-treated with 500 μM HC or DMSO (carrier control) for six hours in crosslinking buffer (X) or non-crosslinking buffer (N). After this pre-treatment step, HC and DMSO were dialyzed out against quenching buffer (20 mM Tris pH 7, 150 mM NaCl) and Lal crosslinking initiated via the addition of crosslinking buffer and incubated for 2 days at 4 ºC. Lal-crosslinked HMWC formation was followed via silver-stained SDS-PAGE and normalized to the untreated, 2-day control (no DMSO or dialysis). Each sample was repeated in duplicate and the normalized cumulative Lal-crosslinked band area for each sample are reported on the right.

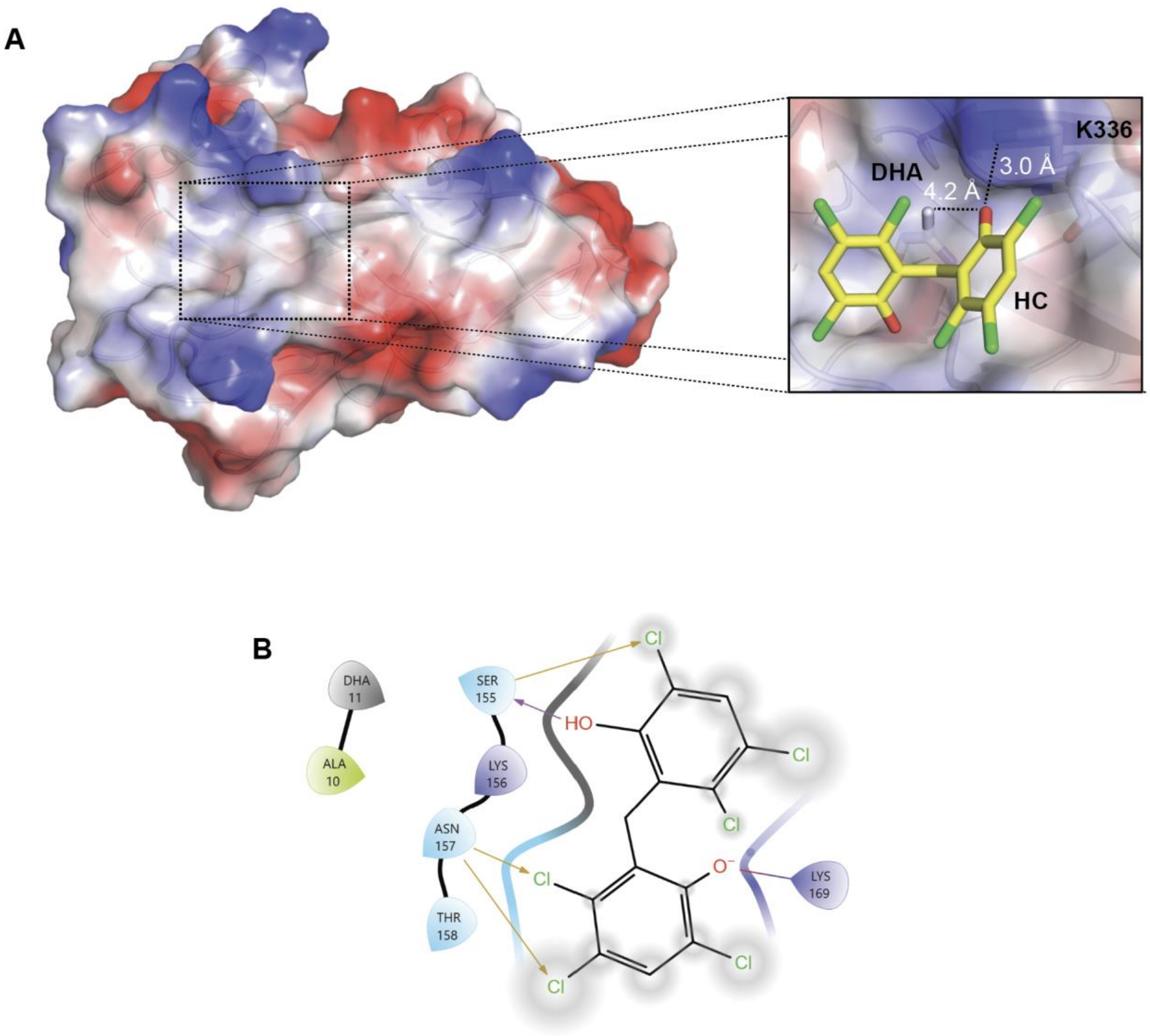


Figure S13: Docking HC to Td FlgE D2 DHA monomer model

**A)** HC binding site of Td FlgE D2$^{DHA}$ monomer model (docking score = -3.883). **B)** Ligand-interaction diagram between Td FlgE D2$^{DHA}$ and HC. Note that Lys169 is equivalent to Lys-336 in the Td FlgE$_{FL}$ dimer model and interaction distance cut-offs were set to 4.0 Å.

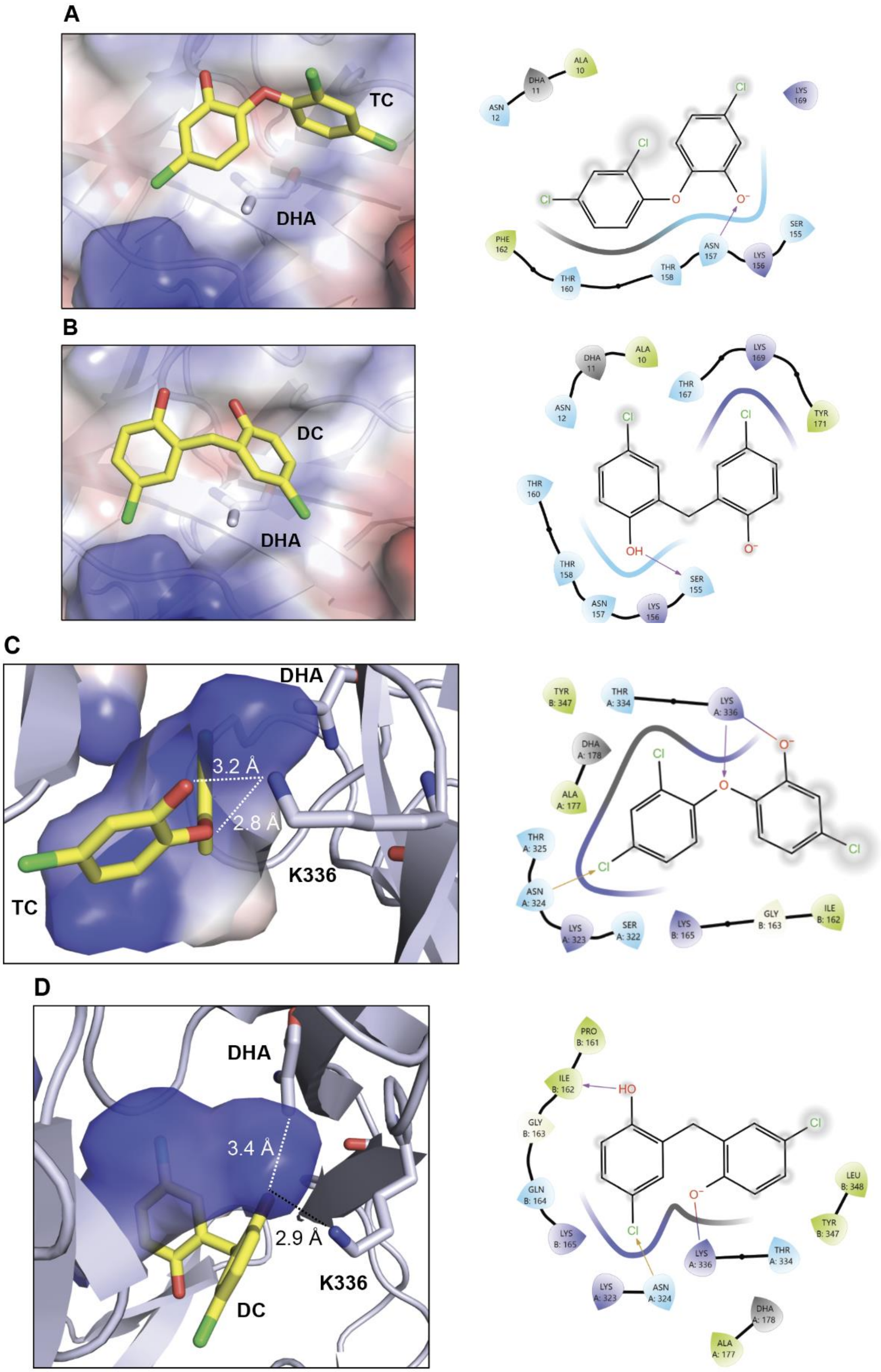
A
TC
DHA
B
DC
DHA
C
DHA
3.2 Å
2.8 Å
K336
TC
D
DHA
3.4 Å
2.9 Å
K336
DC

Figure S14: Top docking poses of TC and DC to FlgE monomer and dimer models.

Top docking pose of (A) TC to our $D2^{DHA}$model, (B) DC to our $D2^{DHA}$ model, (C) TC to our mixed $FlgE_{FL}$ dimer model and (D) DC to our mixed $FlgE_{FL}$ dimer model. For each docking pose, the left panel shows the orientation of TC or DC at the DHA binding site and the right panel shows the ligand interaction diagram for each inhibitor. Ligand interaction distances were cut off at 4.0Å. Residue numbering differences between the $D2^{DHA}$ model and $FlgE_{FL}$ model are as described in Figure S13.

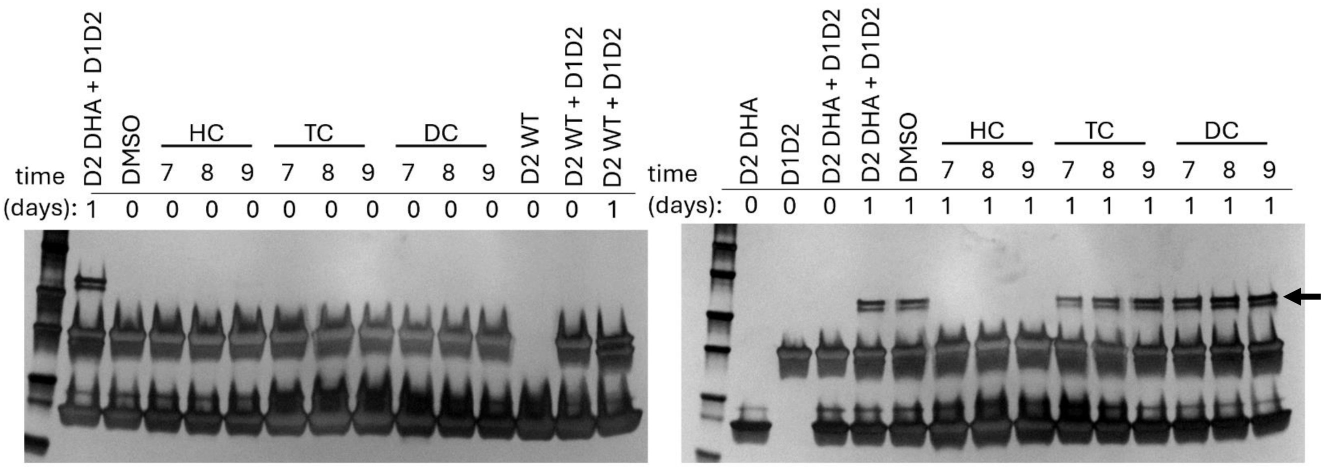


Figure S15: Lal inhibition by HC, TC and DC as a function of pH

100 μM FlgE $D2^{DHA}$ was pre-treated with 500 μM HC, TC and DC for 24 hours at 4°C at a buffer pH of 7, 8 or 9. Each sample was then dialyzed to remove unbound inhibitor and mixed in a 1:5 stoichiometric ratio with D1D2 (D1D2:D2) to initiate Lal crosslinking. Samples were monitored over time (0- and 1-day incubation timepoints) and compared to DMSO-only carrier controls and $D2^{WT}$ negative controls. The presence or absence of the Lal-crosslinked D1D2:D2 heterodimer (indicated with black arrow) was monitored via silver-stained SDS-PAGE.
